# Endogenous Nodal switches Wnt interpretation from posteriorization to germ layer differentiation in geometrically constrained human pluripotent cells

**DOI:** 10.1101/2024.03.13.584912

**Authors:** Miguel Angel Ortiz-Salazar, Elena Camacho-Aguilar, Aryeh Warmflash

## Abstract

The Wnt pathway is essential for inducing the primitive streak, the precursor of the mesendoderm, as well as setting anterior-posterior coordinates. How Wnt coordinates these diverse activities remains incompletely understood. Here, we show that in Wnt-treated human pluripotent cells, endogenous Nodal signaling is a crucial switch between posteriorizing and primitive streak-including activities. While treatment with Wnt posteriorizes cells in standard culture, in micropatterned colonies, higher levels of endogenously induced Nodal signaling combine with exogenous Wnt to drive endoderm differentiation. Inhibition of Nodal signaling restores dose-dependent posteriorization by Wnt. In the absence of Nodal inhibition, micropatterned colonies undergo spontaneous, elaborate morphogenesis concomitant with endoderm differentiation even in the absence of added extracellular matrix proteins like Matrigel. Our study shows how Wnt and Nodal combinatorially coordinate germ layer differentiation with AP patterning and establishes a system to study a natural self-organizing morphogenetic event in *in vitro* culture.

## Introduction

The canonical Wnt pathway is a highly conserved signaling cascade essential to morphogenesis and tissue patterning throughout development. Along with the FGF and TGF-ß superfamily signaling pathways, it is responsible for the formation of the primitive streak (PS), and, subsequently, the mesoderm and definitive endoderm. Perturbation to any of these signals leads to severe gastrulation defects and embryonic lethality (Takada et al., 1994; Conlon et al., 1994; Winnier et al., 1995; Yoshikawa et al., 1997; Liu et al., 1999; Ciruna and Rossant, 2001; Tortelote et al., 2012). Starting at the end of gastrulation, posterior progenitors in the tailbud of the embryo, referred to as neuromesodermal progenitors (NMPs) form the spinal cord and somites in an ordered sequence from anterior to posterior. Continuous Wnt and Fgf signaling is essential for both the formation and maintenance of the NMPs. Disrupting Wnt at this stage results in embryo truncation, failed somite formation, and ectopic neural tubes (Takada et al., 1994; Chapman et al., 1996; Saga et al., 1997; Yoshikawa et al., 1997; Chapman and Papaioannou, 1998; Yamaguchi et al., 1999; Ciruna and Rossant, 2001; P. H. White 2003; Perantoni et al., 2005; Phillip H. White, Farkas, and Chapman, 2005; Olivera-Martinez and Storey, 2007; Dunty et al., 2008). However, it remains unclear how these activities are coordinated and what determines whether Wnt induces primitive streak fates or posteriorizes to give rise to NMPs and their subsequent posterior derivatives.

Quantitative studies of signaling and fate in standard culture are challenging as the colony geometry and cell density are not controlled and can play an important in the response to signals. Previous work has shown that constraining hPSC colonies into circular geometries using micropatterning leads to self-organized germ layer patterning when treated with BMP4 or generates primitive streak-like structures at the colony edge when treated with WNT. In both cases, these self-organized patterns are highly reproducible, providing a robust system for studying signaling dynamics and cell fate patterning *in vitro* (Warmflash et al., 2014; Denglicerti et al., 2016; Martyn et al., 2018; Heemskerk et al., 2019; Chhabra et al., 2019; Martyn et al., 2019A; Martyn et al., 2019B; Minn et al., 2020). Recently, micropatterned differentiation protocols have been developed that generate PSM fates as well as notochord progenitors (Rito et al., 2023; García et al., 2023); however, the mechanisms and signaling dynamics that underlie these fate choices, and particularly how WNT coordinates specifying axial position with germ layer differentiation remains to be elucidated.

In this study, we aimed to develop micropatterned models to study the WNT signaling dynamics leading to posterior progenitors and their derivatives. We confirmed that supplying WNT and FGF signals in standard culture led to NMP-like cells. However, surprisingly, performing the identical protocol in micropatterned colonies led to completely different cell fates. Micropatterned colonies treated with WNT and FGF ligands developed an intricate three-dimensional structure composed of layers of definitive endoderm (DE) cells surrounding an epiblast disk-like cell population. Increasing WNT doses did not shift cell fates towards mesoderm or posterior identity. We show that NODAL, a ligand of the TGF-ß pathway, is upregulated in micropatterns and is responsible for inducing DE cell fates. By measuring signaling and fates, we show that NODAL signaling acts as a switch, affecting the interpretation of WNT and, consequently, cell fate induction. When NODAL signaling is present, WNT is interpreted as a signal to induce primitive steak (PS) fates and subsequent derivatives, which, in the absence of further signaling modulations, leads to anterior endoderm fates. In contrast, when WNT is combined with NODAL inhibition, WNT induces posterior fates in a concentration-dependent manner. Finally, we determined that CHIR, a commonly used WNT activator, induces qualitatively different WNT and NODAL signaling dynamics in this protocol, allowing it to induce posterior mesoderm fates in a dose-dependent manner without requiring additional Nodal inhibition.

## Results

### Geometric constraint changes the outcome of WNT3A and FGF8 mediated differentiation

We set out to create a system to study the WNT signaling dynamics leading to the NMP state and subsequent posterior fates, and how these dynamics differ from those previously reported in giving rise to PS and diverse mesodermal fates (Martyn et al., 2019). NMPs require WNT signaling for both their specification and maintenance. Established protocols have shown that this state can be achieved *in vitro* via activation of WNT (using CHIR99021) and FGF (using bFGF or FGF8) signaling. Using these protocols in standard culture, we observed induction of the NMP state, as measured by the co-expression of SOX2 and BRA (Figure S1A), in greater than 90% of cells. In order to achieve dynamics closer to those that might occur during induction *in vivo*, we substituted WNT3A for CHIR in the protocol that utilizes FGF8. We found that this protocol generated a high fraction of cells that coexpress SOX2 and BRA, albeit lower than with CHIR and bFGF (71 vs 95% co-expression; Figure 1A-D). This shows that treatment with FGF8 and WNT3A ligands in standard culture induces an NMP state in hPSCs.

**Figure 1.**
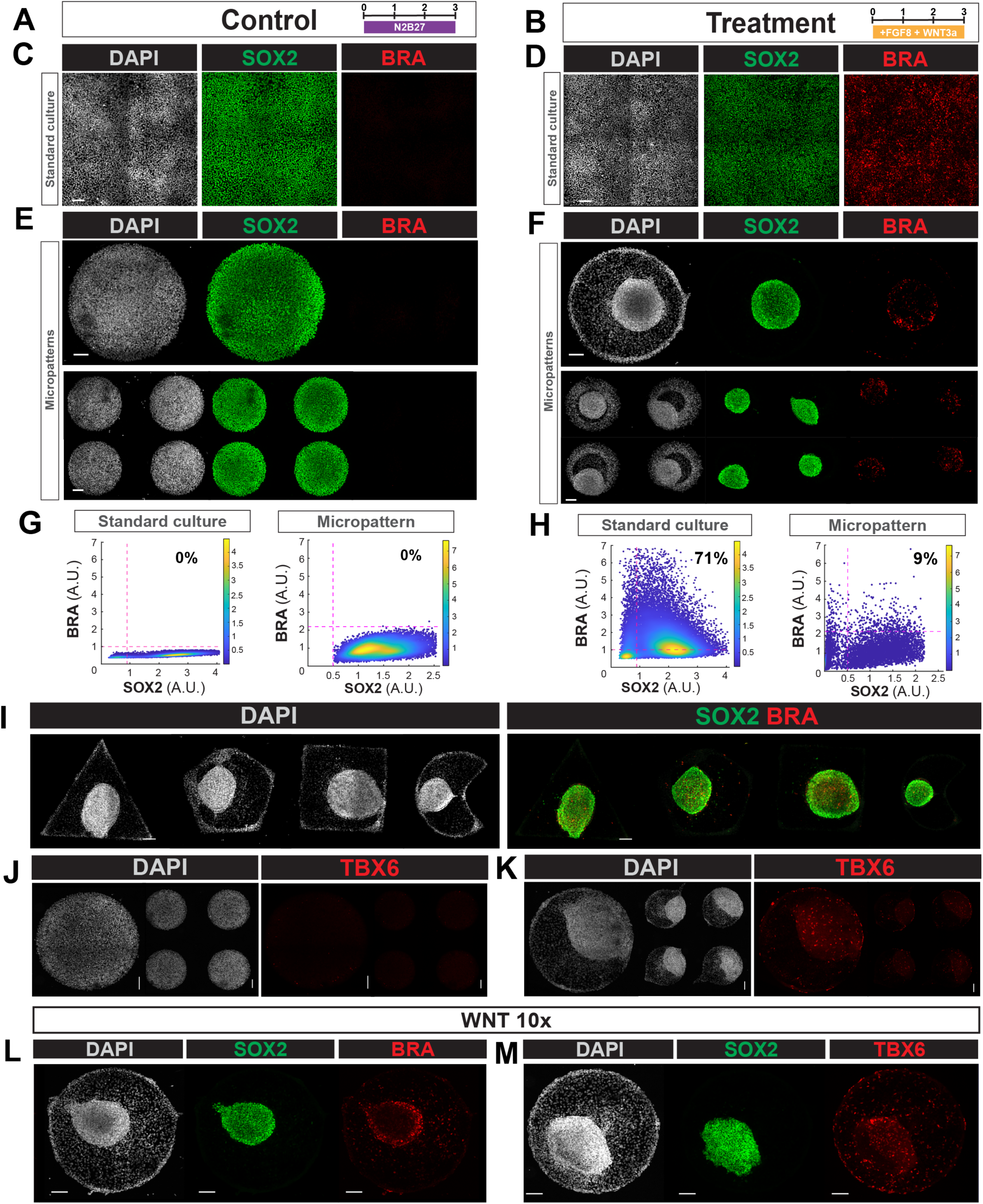
Geometric constraint changes the outcome of WNT3a and FGF8 mediated differentiation. (A, B) Schematics of control (A) and treatment (B) conditions. Concentrations are FGF8 (200 ng/ml) and WNT3a (100 ng/ml). (C, D) Representative immunofluorescent image of the control and treated conditions in standard culture. (E, F) Representative immunofluorescent image of micropatterned colonies in control (E) or treated (F) conditions. Top row 800 µm colonies, bottom row 500 µm colonies. (G, H) Scatterplots of mean intensity (A.U.) for SOX2 and BRA for control (G) and treated conditions (H) in both standard and micropatterned culture. Each dot represents a single cell quantified. Red lines represent the 99th-percent quantiles of the distributions obtained from the negative controls for SOX2 and BRA. Color bar represents the probability density estimate 1×10^−5^. (I) Representative immunofluorescent images of treated micropatterned colonies of different geometries. (J, K) Representative immunofluorescent images of a 800 µm and 500 µm control (J) or treated (K) micropatterned colonies stained for TBX6. (L, M) Representative immunofluorescent image of 800µm micropatterned colonies treated with ten times more Wnt (1000 ng/ml) and stained as indicated. All scale bars are 100 µm. All images are max projections.

We next performed an identical protocol using cells on micropatterns to better study the signaling in space and time that leads to NMPs and subsequent posterior fates. Surprisingly, treatment of micropatterned colonies with FGF8 (200 ng/ml) and WNT3A (100 ng/ ml) resulted in the formation of asymmetric colonies containing a circular cluster of SOX2-positive cells which was offset from the colony center and surrounded by SOX2-negative cells. In both populations, there were a limited number of BRA-expressing cells (Figure 1F,H). A similar pattern emerged on micropatterned colony sizes of 250, 500, 800, and 1000 µm (Figure 1F, Figure S1B). Thus, physical confinement of the colony not only generated spatial ordering but drastically altered cell fate decisions, so that a protocol that generates a high fraction of NMPs in standard culture generates almost none when performed on confined colonies. We hypothesized that the circular geometry of the SOX2-positive cluster might reflect the circular shape of the micropatterns used. To test this, we repeated our treatment on different-shaped patterns, including squares, triangles, and pac-man shapes. Remarkably, the disk-like shape of the cluster of SOX2 cells persisted (Figure 1I), indicating that the SOX2 positive cells self-organize into a circular cluster regardless of the shape of the colony.

We tested whether we observed limited cells with an NMP expression profile because cells had transited through this phase earlier and had already differentiated to more mature cell types by the third day (Figure S1B-C). Indeed, we observed a substantial co-expression of BRA and SOX2 on both day 1 and day 2 of treatment. On day 2, cells at the edge of the colony began losing SOX2 expression, while those at the center maintained their expression, suggesting the initial formation of the SOX2 cluster that is observed on day 3 (Figure S1C). Unlike in standard culture, micropatterned colonies do not maintain co-expression of SOX2 and BRA on day 3, potentially indicating more rapid differentiation.

Previous studies have shown that as NMPs differentiate to pre-somitic mesoderm (PSM), they down-regulate expression of SOX2 and BRA and upregulate the PSM marker TBX6 (Gouti et al., 2014; Lippmann et al., 2015; Wymeersch et al., 2016; Amin et al., 2016; Concepcion et al., 2017). However, we observed only scattered expression of TBX6 in the cells which had downregulated SOX2 and BRA (Figure 1J-K), indicating that the double positive cells that appear on days 1-2 are unlikely to be NMPs but rather cells differentiating to other mesendodermal fates that have not yet downregulated SOX2.

Numerous studies have established a correlation between Wnt levels and induction to mesodermal fates (Chapman et al., 1996; Saga et al., 1997; Yoshikawa et al., 1997; Chapman and Papaioannou, 1998; Yamaguchi et al., 1999; Takada et al., 1994; White, Farkas, and Chapman, 2005; Olivera-Martinez and Storey, 2007; Dunty et al., 2008; Chalamalasetty et al., 2014; Gouti et al., 2014; Lippmann et al., 2015; Wymeersch et al., 2016; Amin et al. 2016; Concepcion et al., 2017), so we tested whether increasing the concentration of WNT would better produce or maintain NMP and PSM cell fates within the colony. Surprisingly, increasing the WNT concentration 10-fold (from 100 ng/ml to 1000 ng/ml) yielded only minor increases in BRA or TBX6 expression, while most of the colony remained negative for these markers as well as for SOX2 (Figure 1L,M). This data indicates that the increase in WNT does not result in a substantial increase of BRA or TBX6 expression. Moreover, in the absence of NMP or PSM differentiation, the identity of both the SOX2-positive and negative cells remained unclear.

### Micropatterned colonies are composed of definitive endoderm and epiblast cells

We utilized RNA sequencing to determine the fates of the cells within the micropatterned colonies. We treated a previously developed SOX2::mCitrine reporter cell line (Camacho-Aguilar et al., 2022) with both Wnt concentrations used above (100 and 1000 ng/ml, referred to as low and high WNT concentrations (WNT 1x and WNT 10x), respectively, from here onwards) and utilized flow cytometry to separate the SOX2-positive cells from the negative ones at the end of treatment (Figure 2A). As expected from imaging, flow cytometry revealed two distinct populations, one which was SOX2 positive and the other SOX2 negative. Increasing the level of WNT decreased the SOX2-positive cell population, leading either to increased SOX2-negative cells or increased cells found in the broad plateau between the two peaks (Figure 2A, S2A).

**Figure 2.**
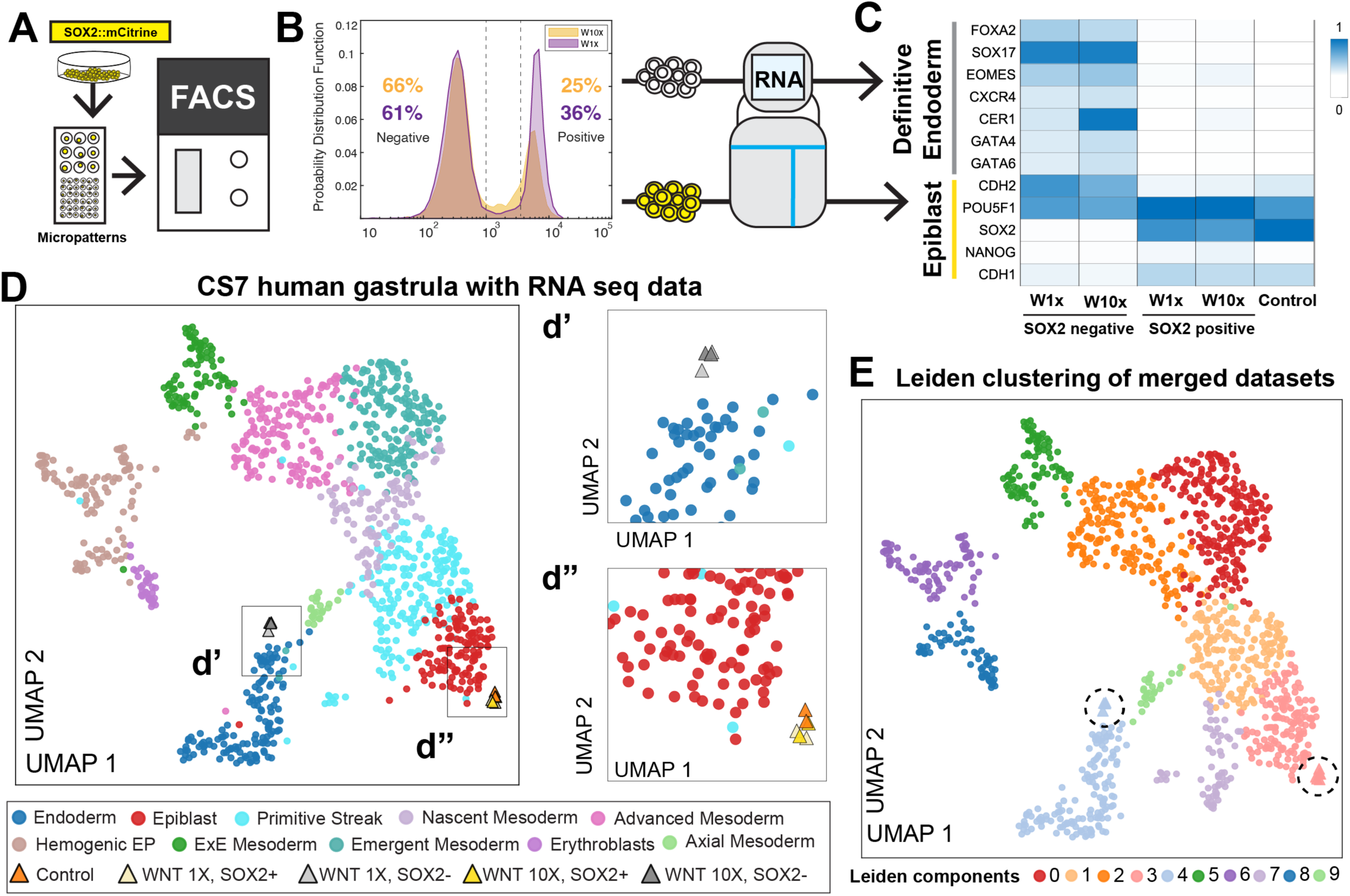
Micropatterned colonies are composed of definitive endoderm and epiblast cells. (A) Schematic illustrating the methodology used to separate cells for RNA sequencing. A SOX2::mCitrine cell line was subjected to treatment, after which flow cytometry was used to separate the SOX2 positive cells from the negative ones. (B) Probability distribution function histogram of SOX2 positive and negative sorted cells for treatments at both WNT concentrations. (C) Heatmap of gene expression within the SOX2 positive (pos) vs SOX2 negative (neg) cells. Expression levels are normalized counts per million. (D) UMAP plot obtained from merging human gastrula CS7 scRNA sequencing dataset in Tyser et al 2021 and our RNA seq data. Cell types are assigned as in Tyser et al 2021. (d’, d’’) Zoomed views from (D). (E) Leiden re-clustering of merged data in (D). Each color represents a different leiden cluster. Circles represent data from Tyser et al. while other shapes represent data from micropatterns as in (D). Dashed circles also highlight the position of micropatterned samples.

Cells separated by SOX2::mCitrine intensity were then subjected to RNA sequencing. Transcriptomic analysis identified the SOX2-negative cell population as definitive endoderm (DE) cells, as indicated by the expression of FOXA2, SOX17, DKK1, CER1, GATA6, OTX2, and N-CAD. Conversely, the SOX2-positive cells displayed other markers of the epiblast, including OCT4, NANOG, and E-CAD (Figure 2C). Both the SOX2 positive and negative cells had very similar transcriptomes at the two different WNT concentrations confirming that the higher levels of WNT shifted the fraction of cells adopting epiblast vs endodermal identities but did not change the fates that are adopted. Through PCA analysis, we found that the first principal component, which represented 70% of the total variation, separated SOX2-negative cells from SOX2-positive cells in treatment and control (i.e. cells grown in N2B27 alone) conditions. The second principal component, which represented 15% of the total variation, separated SOX2-positive cells in the WNT-treated conditions from those in control, untreated replicates (Figure S2B,C). Biological replicates showed a consistent transcriptomic profile.

We then asked whether the cells in treated micropatterns have similar transcriptomes to the corresponding cell types in the CS7 human gastrula dataset from Tyser et al., 2021. We merged the datasets using the scvi-tools package in Python, considering genes that were expressed in both datasets (see Methods section for details) (Lopez et al., 2018; Gayoso et al., 2022). Clustering the cells using Leiden clustering showed that, indeed, the SOX2 positive and negative cells clustered with the *in vivo* epiblast and endoderm populations (Figure 3D,E), respectively, and that these clusters expressed high levels of pluripotency or definitive endoderm marker genes (Figure S2D). In summary, this analysis reveals that micropatterned colonies treated with WNT3A and FGF8 develop into a structure containing an epiblast disk-like shape and surrounding DE cells, and that these populations closely resemble human epiblast and endoderm cells, respectively.

**Figure 3.**
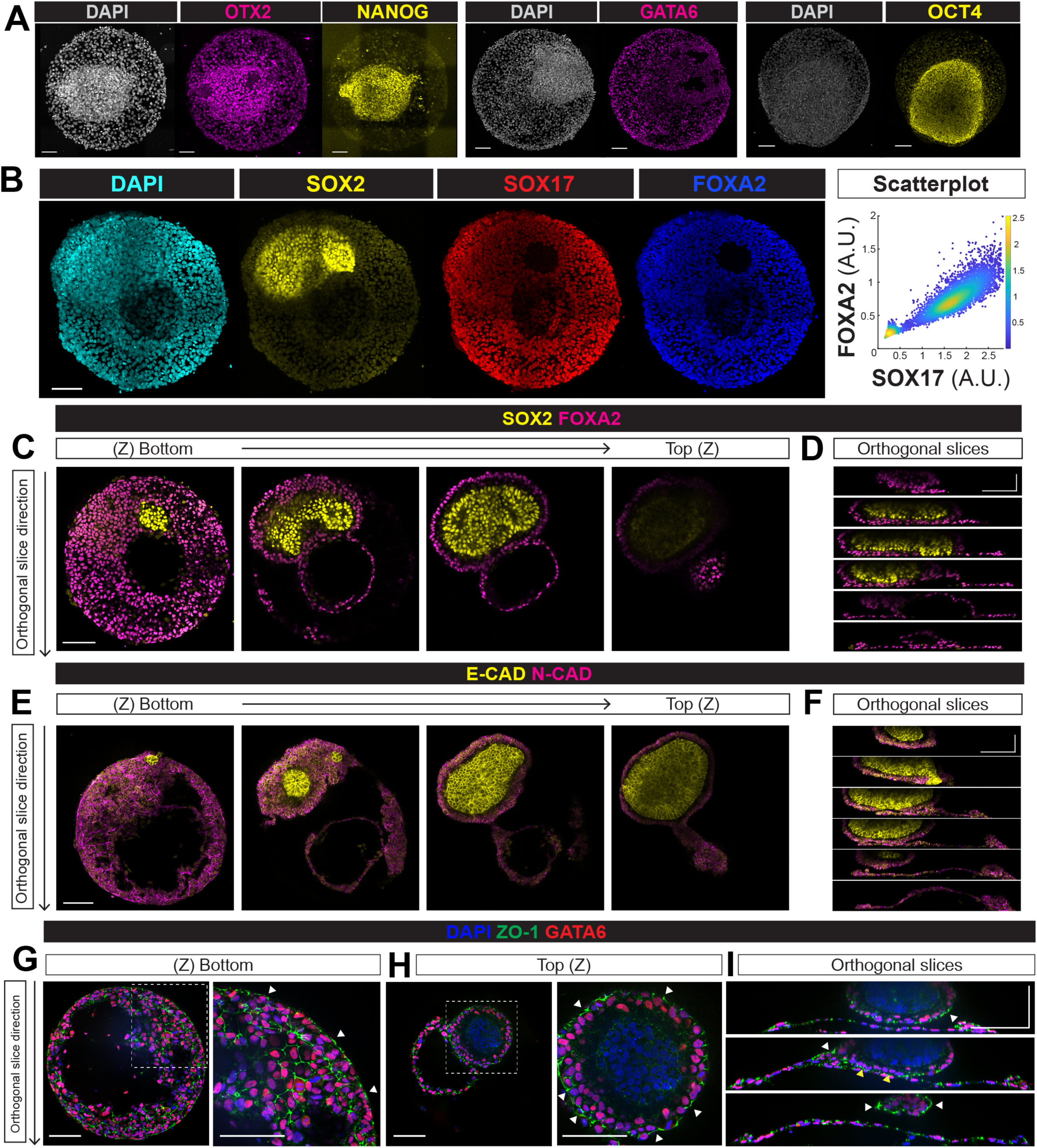
Treated colonies undergo dramatic morphogenesis generating an intricate structure. (A, B) Max projections of representative immunofluorescent images of 700 µm (A) or 500 µm (B) micropatterned colonies treated with WNT 1x with immunofluorescence performed for the indicated proteins. (B,Right) Scatterplot quantifying of mean intensity (A.U.) for SOX17 and FOXA2. Each dot represents a single cell quantified. Color bar is the probability density estimate 1×10^−6^. (C) Individual Z-slices from the same colony as (B). Images shown from left to right correspond from bottom z-plane on the culture surface to the top z-plane.(D) Orthogonal slices from the same treated colony (B). (E, F) Individual Z-slices (E) and orthogonal slices (F) of a colony stained for E-CAD and N-CAD. (G, H) Representative immunofluorescent image of a slice at the bottom (G) or top (H) of a treated colony stained for ZO-1 and GATA6. To the right of each image in a zoomed image of the region indicated in the left panel. White arrows indicate ZO-1 expression at the periphery of the colony. (I) Orthogonal slices from the same treated colony (G,H). White arrows indicate ZO-1 expression facing to the outside of the colony. Yellow arrows indicate inward facing ZO-1 expression between two layers of definitive endoderm. All scale bars are 100 µm.

### Treated colonies undergo dramatic morphogenesis generating an intricate structure

To further validate the transcriptomic results, we immunostained colonies with multiple markers of epiblast and endoderm fates. Our results show that, indeed, the disk-like cell population expressed SOX2, OCT4, NANOG, and E-CAD, while the remaining portion of the colony displayed expression of FOXA2, SOX17, GATA6, OTX2, and N-CAD (Figure 3A-B). The expression of OTX2 confirmed the anterior identity of the DE cell population. Interestingly, while previous studies have shown the suppression of anterior fates through elevating Wnt signaling (Gouti et al., 2014; Tsakiridis et al. 2014; Lippmann et al. 2015; Gouti et al., 2017; Metzis et al. 2018; Frith et al. 2019), OTX2 expression remained unaffected in our colonies when we increased WNT levels. This indicates that increasing WNT signaling does not result in the induction of posterior identities in this context.

High-resolution imaging in three dimensions showed that colonies were composed of a layer of definitive endoderm (DE) cells at the base, extending across the entirety of the colony. At the colony edges, this layer appeared to fold over so that a portion of the colony contained a double layer of endoderm at the bottom (Figure 3C,D). The endoderm underneath the epiblast population was always double-layered in this manner and curved upwards, forming a cup around the epiblast cluster. In many colonies, a narrow “stalk” of cells extended off of the epiblast population and remained adhered to the surface (Figure 3C-F), while in others, the endoderm completely displaced the epiblast cells from the culture surface (Figure 3G-I). In nearly all colonies, a “bubble” consisting of DE cells that detached from the surface appeared next to the cup (Movie S1). Expression of E-cadherin (E-CAD) and N-cadherin (N-CAD) within the colony was mutually exclusive, with E-CAD expression localized to the epiblast disk-like cells, whereas N-CAD was associated with the DE cells (Figure 3E,F).

Next, we stained for ZO-1, which marks apical tight junctions, to investigate if there was an acquisition of a basal-apical polarity. Where the endoderm was single-layered, we observed ZO-1 on the top of the layer facing the media and pointing to the outside of the colony at the edge (Figure 3G). The DE cells surrounding the epiblast disk-like cells displayed a similar expression pattern of ZO-1, with the apical side facing outward and the basal side facing the epiblast-like cells (Figure 3H). In places where the endodermal layer was double-layered, typically both layers pointed their apical layers inward so that they faced each other, with both basal sides facing outward in opposite directions (Figure 3I).

To observe the formation of the epiblast disk-like cell population, we employed live-cell confocal microscopy to monitor colonies of SOX2::mCitrine and E-CAD::mCitrine cells (Liu et al., 2022; Camacho-Aguilar et al., 2022) during the second and third day of treatment. We observed initially, SOX2 is expressed throughout the entire colony, but then it is downregulated at the edge. On the third day, these SOX2 negative cells undergo large-scale movements toward the colony center. These movements displace the SOX2-positive cells inwards into a cluster and lift them off the surface. Colonies treated with higher levels of WNT underwent a similar sequence of events, albeit slightly accelerated in time (Movie S2) (Figure S3). E-CAD expression showed essentially identical dynamics to SOX2, indicating that it is the same population of cells that expresses both of these markers (Movie S3) (Figure S3). Consistently, immunostaining confirmed the expression of other epiblast markers in the E-CAD positive population (Figure S3). In contrast, untreated colonies maintained SOX2 and E-CAD expression throughout and did not develop the same complex morphology. Performing an identical protocol on two additional hPSC lines, H9 and RUES2, yielded consistent results, including the presence of self-organized colonies with distinct epiblast and endodermal cell populations (Figure S2). In summary, these data indicate that exposure to FGF8 and WNT3A in micropatterns, unlike in standard culture, leads to dramatic morphogenesis concomitant with endoderm differentiation.

### ß-catenin signaling dynamics reveal a dose-dependent response to WNT

We next examined the dynamics of WNT signal transduction using a GFP::ß-catenin reporter cell line which also expresses H2B::RFP for nuclear identification and tracking (Massey et al., 2019). ß-catenin is the central signal transducer for the WNT pathway, and activation of the pathway results in inhibition of the destruction complex causing accumulation of ß-catenin in the cell nucleus where it activates transcription. Our RNAseq data above indicated largely identical cell fates at both low and high doses of WNT. Immunofluorescence confirmed virtually identical expression patterns of DE and epiblast disk markers as well as the same elaborate morphogenesis at both concentrations of WNT (Figure 4A,B). Thus we measured ß-catenin dynamics at both low and high doses of WNT to see whether these doses were translated into distinct ß-catenin responses. We segmented individual nuclei in three dimensions using the H2B::RFP channel and quantified the mean ß-cat intensity over the nuclear volume for all nuclei in each colony (Figure 4D).

**Figure 4.**
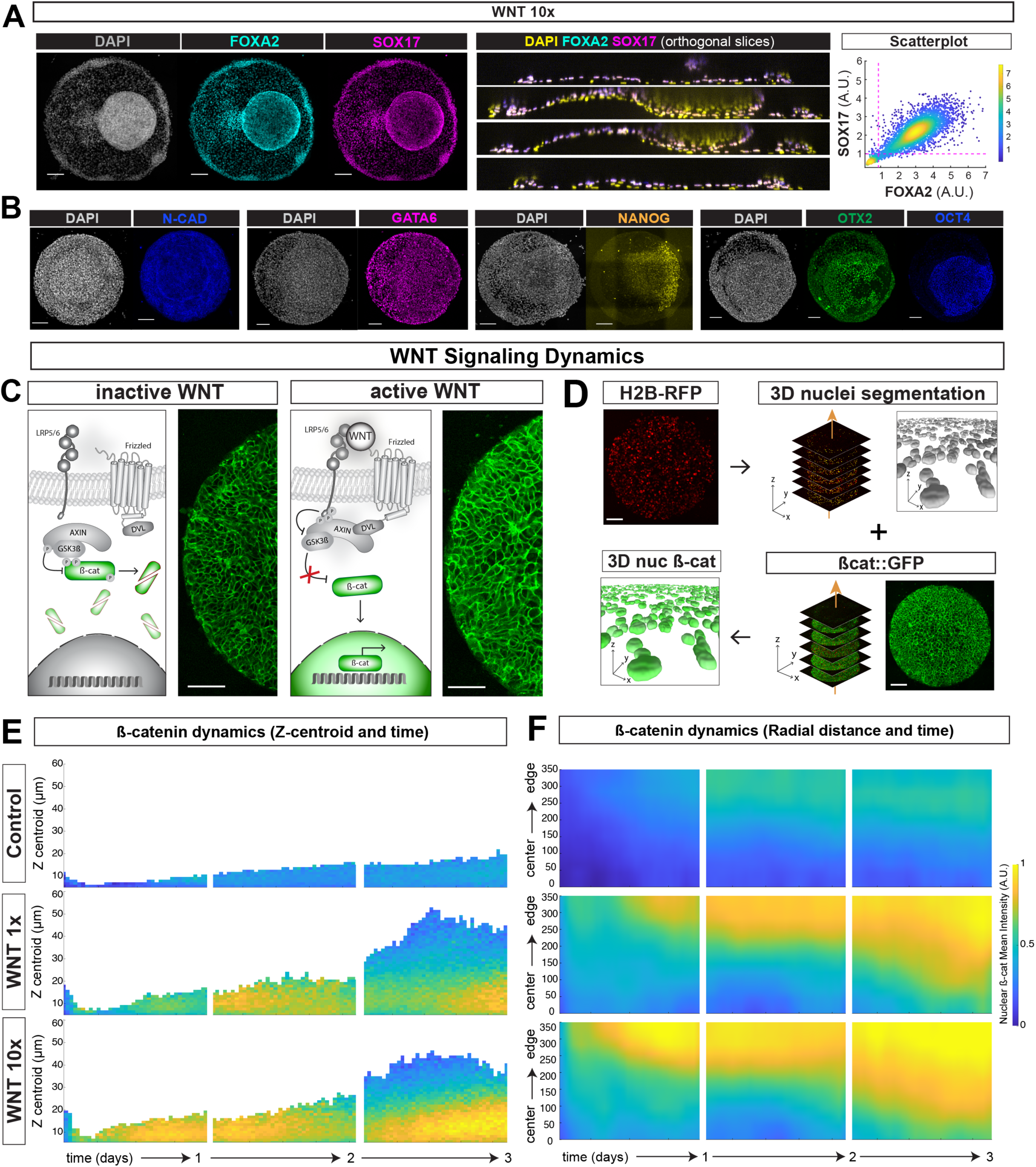
ß-catenin signaling dynamics reveal a dose-dependent response to WNT. (A) Representative immunofluorescence image of 800 µm colony treated with WNT 10X and stained as indicated. Left, Max projection. Middle, orthogonal slices of the same colony. Right, Scatterplot quantifying single cell expression for FOXA2 and SOX17. Red lines represent the 99th-percent quantiles of the distributions calculated from the negative controls. Color bar represents the probability density estimate. (B) Representative immunofluorescence max projection images of 700 µm colonies treated with WNT 10x. All scale bars are 100 µm. (C) Schematics of the Wnt pathway schematic and images of the GFP::ß-cat reporter cell line with (right) or without (left) WNT activity. (D) Workflow for quantification of nuclear ß-catenin. Nuclei segmentation was performed on the H2B::RFP z-stack at each timepoint. The resulting nuclear mask was used to quantify GFP::ß-cat mean intensity levels for each cell. (E) Kymograph quantifying average nuclear ß-cat levels as a function of the Z-centroid of the cells and time. (F) Kymograph quantifying average nuclear ßcat levels as a function the distance from the center of the colony and time. In (E,F), each colored square represents the mean intensity of nuclear ßcat levels of all cells quantified at that position during that time. Live imaging was performed for 22 hours each day.

Control colonies in N2B27 that were not treated with WNT maintained low levels of nuclear ß-catenin throughout, with only a slight increase in ß-catenin levels toward the end of the three-day culture period likely due to endogenous WNT signaling between cells at high density. These control colonies remained relatively flat and did not undergo morphogenesis (Fig 4E-F Control).

When treated with the low dose of WNT (WNT 1x), cells at the colony edge initially responded and reached high levels of signaling by approximately 16h of treatment (Fig 4F). The region of active signaling spread inwards towards the end of the first day and then stabilized with a region of high signaling that maintained a fixed width over the course of the second day of treatment (Fig 4F). Finally, during the third day, signaling appeared to spread inward again, this time more rapidly, which corresponded to the morphogenesis of the colony described above (Fig 4F). As the cells differentiating to DE moved under the remaining epiblast cells, the region of WNT signaling expanded, reflecting the spread of these cells with high signaling levels. Signaling in the remaining epiblast cells remained low, as can be seen by viewing the dynamics as a function of z coordinate and time (Fig 4E). As the DE cells moved toward the center of the colony expanding the WNT signaling inward, the epiblast cells are pushed to higher z-coordinates and maintain low WNT signaling. Thus, differentiation initially involves WNT signaling, which fills a territory of fixed width and causes definitive endoderm differentiation. These cells then spread and drive a morphogenetic event during which the epiblast cells are pushed up into a ball and retain low levels of WNT signaling. Colonies treated with the higher WNT concentrations (WNT 10x) showed qualitatively similar dynamics with higher levels of nuclear ß-cat compared to the WNT 1x colonies across time. In both WNT conditions, the dynamics of WNT signaling are consistent with that of the downregulation of SOX2 and E-CAD (Figure S3). Thus, cells do show a stronger response to the higher dose of WNT despite differentiating to an identical set of cell fates.

### NODAL drives definitive endoderm cell fates while restricting posterior mesoderm differentiation

The differentiation of cells to definitive endoderm using only WNT and FGF on micropatterns is surprising as previous studies indicated that TGF-ß/Activin/Nodal signaling (hereafter Activin/Nodal signaling) is essential for generating such cell fates *in vitro* (D’Amour et al. 2005; Yasunaga et al. 2005; Spence et al. 2011; Toivonen et al. 2013; Jacob et al. 2017; Miller et al. 2019; Daoud and Múnera 2020). Therefore, we hypothesized that an endogenous ligand could be activating this pathway to induce DE cell fates. Levels of this ligand could represent an important difference between standard and micropatterned cultures. To test this, we treated the colonies with SB431542 (SB), an inhibitor of the ACTIVIN/NODAL receptors ALK4/5/7 (Figure 5A). Our results showed that inhibition of ACTIVIN/NODAL signaling suppressed DE cell fates, and no expression of FOXA2 or SOX17 was detected (Figure 5B). In contrast, inhibition of BMP signaling by LDN193189 (LDN), did not affect DE differentiation at either WNT concentration (Figure S5 D,E). When both inhibitors were used (also known as dual SMAD inhibition), we observed no expression of SOX17 and scarce FOXA2 consistent with the results obtained with the ACTIVIN/NODAL inhibitor alone (Figure S5 F,G). These results show that endogenous signaling through the Activin/Nodal branch of the TGF-ß pathway is responsible for DE cell fates (Figure 5B).

**Figure 5.**
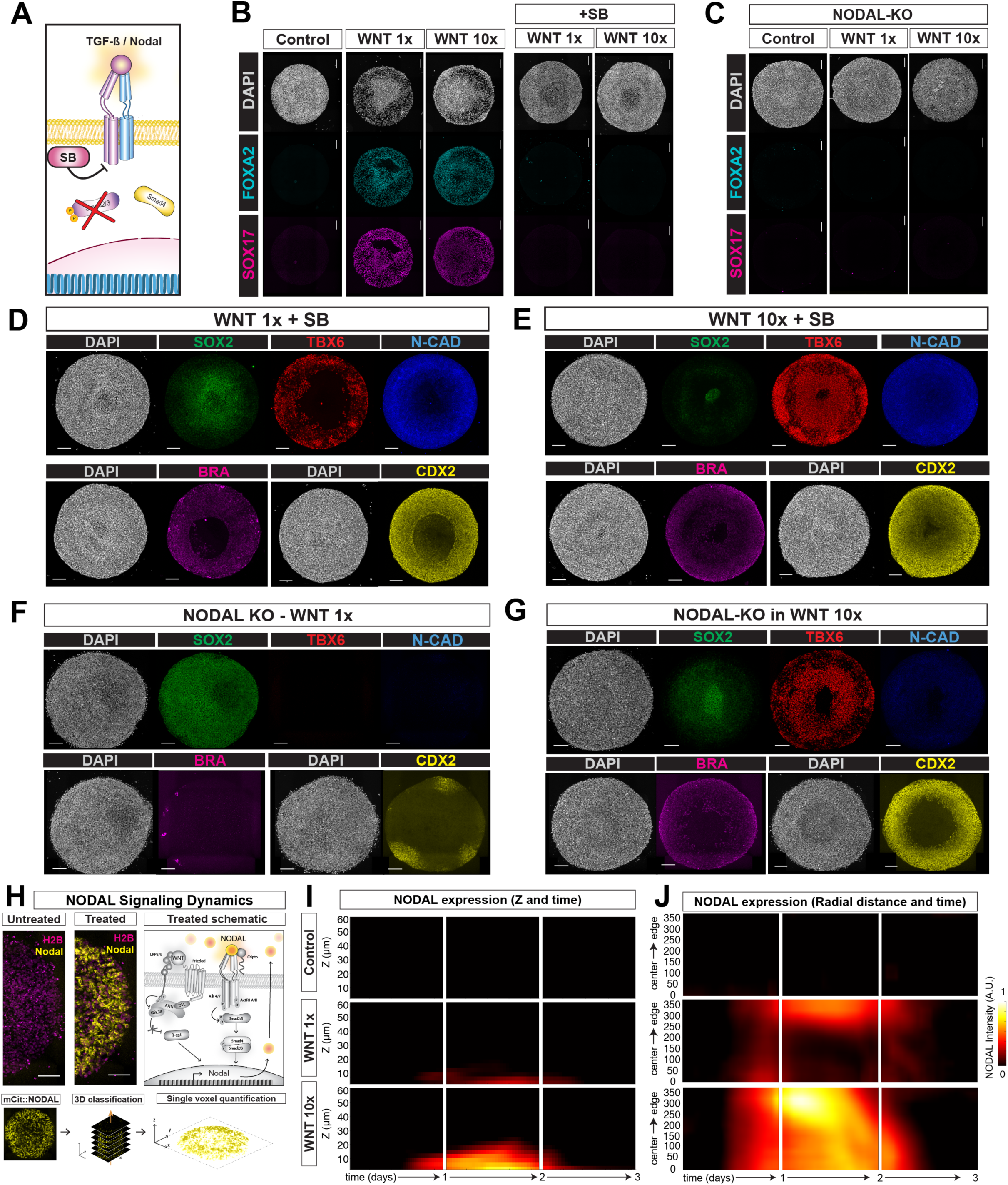
NODAL drives definitive endoderm cell fates while restricting posterior mesoderm differentiation. (A) Schematic of the TGF-ß/ACTIVIN/NODAL pathway inhibition and the action of the receptor inhibitor SB. (B) Representative immunofluorescent images showing the effect of SB treatment (B) or NODAL knockout (C) on endoderm markers SOX17 and FOXA2. (D,E) Representative immunofluorescent images of micropatterned colonies treated with the indicated WNT concentration and the TGFb inhibitor SB. (F,G) Representative immunofluorescent images of micropatterned colonies of NODAL knock-out cells treated with the indicated WNT concentration. All images are max projections. All scale bars are 100 µm. (H) Schematic of the Nodal pathway and representative images of the NODAL::citrine reporter in the active and inactive states. Bottom, Workflow diagram for quantifying NODAL expression. (I) Kymograph quantifying NODAL expression as a function of Z and. time. (J) Kymograph quantifying NODAL expression as a function of distance from the center of the colony and time. In (I,J), each colored square represents the mean intensity of NODAL::citrine levels of all cells quantified at that position during that time.

We tested whether inhibition of ACTIVIN/NODAL diverted cells that would have adopted DE fates to NMP and PSM fates. Indeed, the addition of SB to our treatment allowed the induction of NMP (SOX2+, BRA+) and PSM (TBX6+, CDX1+, CDX2+) cell types (Figure 5D). Remarkably, when combined with SB treatment, increasing WNT had a significant effect as it resulted in colonies consisting almost entirely of PSM cell fates (Figure 5E). Thus, cell fates are sensitive to WNT signaling levels, but only when ACTIVIN/NODAL signaling is inhibited. Moreover, dual-SMAD inhibition led to the formation of colonies with similar patterns, with posterior and PSM cell fates acquired, while DE cell fates were absent (Figure S5F,G). Under dual-SMAD inhibition, we observed expression of SOX1 and SOX2 in the center, indicating neural progenitors derived from the NMP state (Figure S5G).

We next asked which ligand of the ACTIVIN/NODAL pathway is responsible for inducing DE and hypothesized that NODAL, known to be required for mesendoderm differentiation during gastrulation *in vivo*, may play this role *in vitro* as well. To test this, we applied the same treatments to a previously developed NODAL knock-out (NODAL-KO) cell line (Chhabra et al 2019). We found no expression of SOX17 or FOXA2 in the NODAL-KO line, indicating that NODAL is responsible for inducing DE cell fates (Figure 5C). While NODAL-KO colonies abolished DE cell fates, they failed to upregulate posterior markers, with only small patches of CDX2 and no TBX6 expression. Instead, SOX2 was expressed throughout the colony. Increasing WNT in the NODAL-KO cell line generated colonies comprised of NMPs and PSM, similar to the expression patterns when NODAL was inhibited with the small molecular inhibitor (Figure 5G). These results provide further evidence that NODAL is involved in the decision between endoderm and mesoderm.

After determining that NODAL was responsible for DE cell fates, we proceeded to study its expression dynamics using a previously reported cell line with a NODAL::mCitrine fusion at the endogenous locus (Liu et al., 2022) (Figure 5H). Untreated colonies did not express detectable levels of NODAL. In contrast, with WNT 1X treatment, we observed that NODAL production started simultaneously in two territories, one at the edge and the other at the center at 12 hours (Figure 5J). NODAL expression subsequently expanded through the entire colony by the end of the first day. NODAL levels then increased at the edge and declined at the center, so that expression was restricted to the edge. On the third day, NODAL expression was again detected in the center before declining and becoming undetectable at approximately 60 hours post-treatment (Figure 5J). NODAL expression was only detectable close to the surface of the culture (Figure 5I), indicating that after the onset of differentiation, it was active only within the DE cell population and never reached the epiblast cells positioned at higher Z-levels. Treatment with higher levels of WNT (WNT 10X) led to higher NODAL expression levels which peaked at the edge and were detectable throughout the colony from late on the first day through the beginning of the third day (Figure 5I-J). In this case, as well, NODAL expression was restricted to the endoderm layer close to the surface and was not found in the overlying epiblast cells (Figure 5I).

Collectively, these results show that WNT signaling triggers NODAL in a dose-dependent manner, and together these signals induce DE cell fates. When NODAL is inhibited, cell fates divert to PSM. Thus, the interpretation of the WNT signal is modulated by the presence of NODAL. With NODAL, cells adopt DE fates in a manner that is largely insensitive to the dose of WNT, while cells adopt posterior fates in a dose-dependent manner when NODAL is inhibited.

### Definitive endoderm and pre-somitic mesoderm divergence depends on the timing of Nodal inhibition

We next investigated the impact of the timing of NODAL signaling on DE cell fate specification by inhibiting NODAL at different time points. First, we added SB from 24 hours until the end of treatment (24-72h) (Figure 6A). Remarkably, in this condition, PSM cell fates dramatically expanded at the expense of DE in both WNT concentrations, even compared to when NODAL is inhibited throughout (Figure 5D-E, 6A). FOXA2 and OTX2 were absent, while TBX6 and CDX2 were expressed throughout the colonies (Figure 6A). Moreover, when increasing WNT, we observed full suppression of SOX2 expression, resulting in a colony with all cells adopting the PSM fate (Fig 6A). These results show that allowing Nodal signaling during the initial 24 hours enhances posterior mesoderm induction.

**Figure 6.**
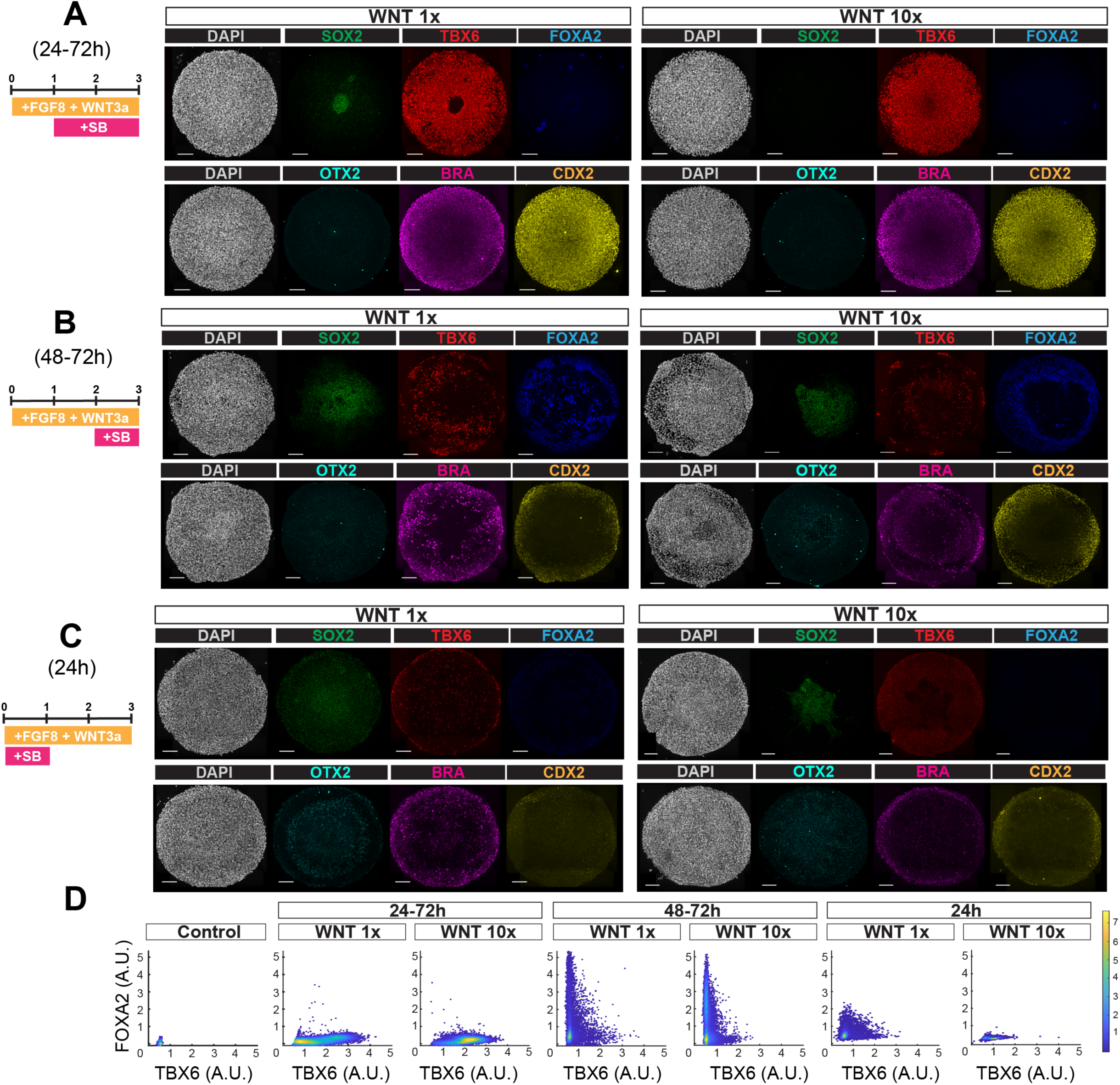
Definitive endoderm and pre-somitic mesoderm divergence depends on the timing of Nodal inhibition. (A - C) Representative immunofluorescent images of micropatterned colonies treated with SB for the indicated time periods and with the indicated WNT concentrations. All scale bars are 100µm. All images are max projections. (D) Scatterplots of FOXA2 vs TBX6. Color bar indicates probability density function 1×10^−5^. Each dot represents a single cell quantified.

Next, we investigated the effect of introducing SB from 48-72h (Figure 6B). We observed scattered expression of PSM and DE cell fates. A center of SOX2-positive cells remained with an outer ring of CDX2. Interestingly, increasing WNT expanded DE cell fates in a broader outer ring that also expressed CDX2 in a thin border region, indicating potential hindgut cell fate (Fig 6B). These results show that, at this time point, some cells already have committed to DE while others retain the ability to shift to PSM fates, resulting in colonies with mixtures of these fates.

Finally, we inhibited NODAL only for the first 24 hours (24h) (Figure 6C). We observed a weaker expression of both DE and PSM markers compared to previous conditions. OTX2 and FOXA2 were only observed under WNT 1x, while BRA and TBX6 cell fates predominated at the border, with SOX2 being expressed in the center of the colony (Figure 6C). This pattern was similar to that observed upon adding SB at 48 hours (48-72h) (Figure 6B). In contrast, increasing WNT resulted in the loss of DE cell fates and an increase of TBX6 positive cells throughout the colony, but its expression was much lower when compared to the other inhibition conditions (Fig 6C). This data indicates that inhibiting the TGF-ß pathway for the first 24 hours negatively impacts differentiation of both endoderm and mesoderm fates, likely by impairing the initial step of primitive streak differentiation. Thus, NODAL is broadly involved in the initial step that leads to diverse mesendodermal fates during the first 24 hours of differentiation, while at later time points, it alters WNT signal interpretation, and is the main determinant of whether cells differentiate to endoderm or posterior mesoderm.

### GSK3ß inhibition bypasses the requirement for NODAL inhibition in inducing posterior cell fates

CHIR99021 is a GSK3ß inhibitor commonly employed as a Wnt activator and is widely employed in diverse differentiation protocols that rely on Wnt signaling (Gouti et al. 2014; Lippmann et al. 2015; Wymeersch et al. 2016; Amin et al. 2016; Concepcion et al. 2017; Gouti et al., 2017; Beccari et al. 2018; Matsuda et al. 2020; Budjan et al. 2022; van den Brink and van Oudenaarden 2021; Xu et al. 2021; Sanaki-Matsumiya et al. 2022; Miao et al. 2023; Rito et al., 2023). This molecule bypasses ligand-receptor dynamics, and we wondered whether colonies treated with CHIR would have different signaling dynamics and pattern differently compared to those treated with WNT3A. At 3µM of CHIR, we found substantial induction of PSM cell fates with an outer ring of DE and low expression of SOX2 in the center (Figure S7A). When adding FGF8 together with CHIR, a similar pattern was observed, but the outer ring of DE increased and the SOX2 center disappeared (Figure S7B). Next, we increased the CHIR dose to 10 µM and found colonies expressing only markers of PSM fates (BRA, TBX6, and CDX2) (Figure 7A). SOX2, OTX2, SOX17, and FOXA2 were completely absent (Figure 7A). Interestingly, neither concentration of CHIR, regardless of whether it was combined with FGF8, resulted in the complex morphogenesis that results from FGF8 and WNT3A treatment. To understand whether the absence of DE cell fates was due to reduced Nodal expression, we monitored expression dynamics and found that CHIR induced NODAL at similar levels to the higher dose of WNT (Fig 7B,C). However, the duration of NODAL expression induced by CHIR was only 24 hours, in contrast to that induced by WNT 10x, which lasted approximately 36 hours (Fig 5I,J). Monitoring WNT signaling dynamics using the ß-catenin reporter cell line, showed that CHIR treatment resulted in drastically elevated WNT signaling levels compared to all other conditions (Figure 7D-F). Thus, CHIR induces both much stronger WNT signaling and more transient NODAL signaling compared to WNT treatment, explaining why cells adopt PSM rather than DE fates. Notably, WNT signaling activity induced by WNT3A was similar whether or not colonies were treated with SB (WNT 10x and WNT 10x+SB), further confirming that with WNT treatment, NODAL expression, rather than WNT signaling activity, is the decisive factor between mesoderm and endoderm cell fates.

**Figure 7.**
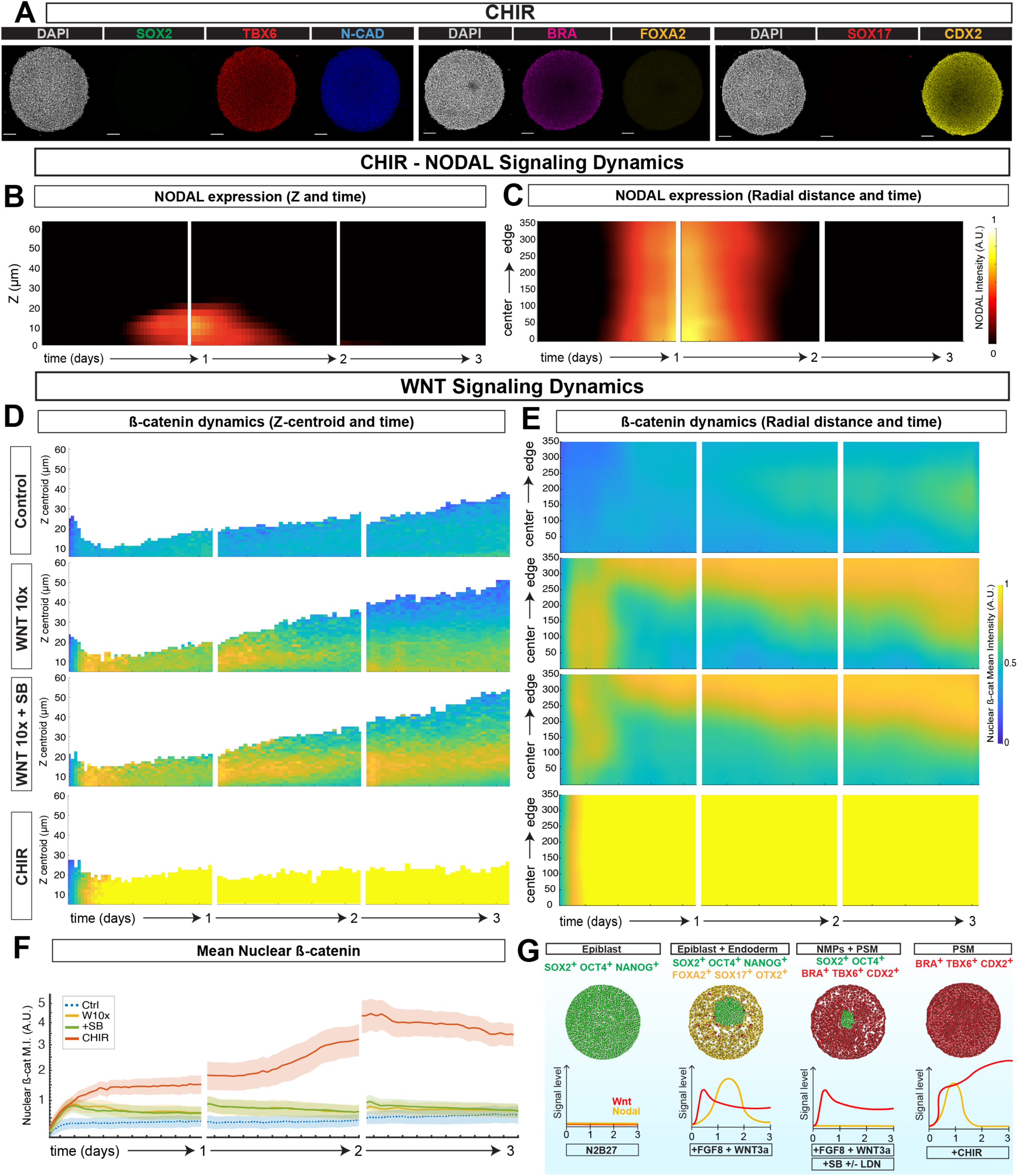
GSK3ß inhibition bypasses the requirement for NODAL inhibition in inducing posterior cell fates. (A) Representative immunofluorescent max projection images of CHIR treated colonies. Scale bars 100µm. (B,C) NODAL signaling dynamics of CHIR colonies. (B) Nodal signaling as a function of Z and time (C) NODAL signaling as a function of distance from the center and time (D, E) WNT signaling dynamics for the indicated treatments as a function of either the Z-centroid and time (D) or distance from the center and time (E). In (B-E) each colored square represents the mean intensity of the reporter for all cells quantified at that position during that time. (F) Mean intensity of nuclear ß-cat of all conditions. (G) Summary Diagram. Nodal determines cell fate specification downstream of WNT ligands but with high concentration of CHIR, Nodal inhibition is expendable.

In conclusion, these findings show that DE is driven by NODAL downstream of WNT. When NODAL is inhibited, cells acquire PSM cell fates in a WNT concentration-dependent manner. However, CHIR is able to induce colonies composed entirely of PSM cells through different signaling dynamics: an acute, more transient production of Nodal and significantly higher Wnt activity levels compared with WNT3A (Figure 7G).

## Discussion

Our results provide evidence that physical constraint impacts cell fate decisions and morphogenesis. While treatment of hPSCs with FGF8 and WNT3A in standard culture induces an NMP-like state, on micropatterns, differentiated colonies are comprised of two spatially segregated populations - epiblast and DE, which undergo dramatic morphogenesis to form a complex 3D structure. In this context, higher levels of WNT stimulation are translated into higher levels of nuclear ß-catenin, but cell fates remain unaffected. This DE differentiation results from endogenous NODAL combined with exogenous WNT, and if NODAL is inhibited, cells adopt posterior mesoderm fates in a concentration-dependent manner. WNT activation via GSK3ß inhibition with CHIR leads to qualitatively different WNT and NODAL dynamics, allowing posterior cell fate differentiation in a concentration-dependent manner without the need for TGF-ß inhibition. Our results collectively demonstrate the intricate interplay between Nodal and Wnt that coordinates axial patterning and germ layer differentiation.

The lack of DE differentiation in response to WNT and FGF in standard culture likely results from reduced NODAL signaling compared to such on micropatterns. In standard culture, the colony area continuously expands and the cell density does not reach a sufficiently high density for endogenous NODAL signaling between cells to reach levels required for inducing DE. In contrast, micropatterned colonies rapidly achieve a higher density in a confined space, leading to higher concentrations of NODAL that, along with the WNT ligand, induce DE cell fates. This difference in signaling dynamics in culture methods aligns with our previous observations with BMP treatment, where NODAL signaling is observed upon BMP4 exposure in micropatterned colonies but not in standard culture (Liu et al., 2022). *In vitro* differentiation protocols with hPSCs have shown that combined treatment with Wnt and Activin, a Nodal substitute, is sufficient to drive DE specification (D’Amour et al., 2005; Spence et al., 2011; Toivonen et al., 2013; Pagliuca et al., 2014; See Loh et al., 2014; Huang et al., 2015; Jacob et al., 2017; Zhang et al., 2017; Broda, McCracken, and Wells, 2019; Miller et al., 2019). Thus, adding Nodal stimulation to NMP protocols in standard culture is sufficient to convert them to DE-producing protocols.

We found that micropatterns treated with WNT and FGF first form a territory of definitive endoderm in a region at the edge of the colony where WNT signaling is high, and that morphogenesis occurs downstream of this differentiation. While the morphogenesis is unique to this model, the patterns of signaling are consistent with those observed in previous studies of micropatterned cultures. In BMP-treated micropatterns, the exogenous BMP signal is primarily received at the edge, similar to the exogenous WNT treatment here (Warmflash et al., 2014; Deglincerti et al., 2016; Morgani et al., 2018; Chhabra et al., 2019; Heemskerk et al., 2019; Jo et al., 2022; Liu et al., 2022). Colonies in pluripotency media treated with WNT or a combination of WNT and Activin induce a border region of mesoderm or endoderm, respectively (Martyn, Brivanlou, and Siggia, 2019A; Martyn, Brivanlou, and Siggia, 2019B). These observations and our results underline the influence of the boundary in driving differentiation as well as the combinatorial influence of WNT and NODAL in mesendodermal cell fate decisions.

Colonies treated with WNT3A and FGF8 undergo morphogenesis which leads to a circular cluster of epiblast cells sitting in a cup of endodermal cells. This cluster is circular regardless of the shape of the colony, indicating self-organization that does not rely on the boundary. Next to this cup, the endoderm makes a bubble above the culture surface with a hollow space underneath. One possible mechanism by which this structure could form is that the entire epiblast portion of the colony may be under tension, which causes it to tend to assume a circular shape. This tension is balanced with adhesion to the culture surface which initially keeps the colony flat. Once the endoderm differentiates, it begins to migrate under the epiblast cells and severs them from the surface. Once a sufficient amount of contact area is lost, the tension within the epiblast compartment rapidly causes it to condense into a ball. Because the epiblast is stuck to the endoderm beneath it, this part is pulled into a cup shape which holds the epiblast ball from below, while the resulting stretching of the endoderm forms the bubble next to the cup. Measuring the distribution of molecules such as myosin, along with methods of measuring or inferring forces, could confirm this hypothesis and yield more insight into this process.

Three-dimensional morphogenesis in cell culture is typically induced by adding an extracellular matrix (ECM) to the culture medium, and the extensive morphogenesis we observed was surprising as ECM proteins are not included in our treatments. However, transcriptomic analysis revealed that the DE population expressed high levels of ECM proteins, including COL-IV-alpha, COL-IX-alpha, and Fibronectin. These proteins are also found in the endoderm of E6.5 mouse embryos (Futaki et al., 2019) and likely contribute to forming the structures we observed. The DE cells also expressed high levels of WNT inhibitors DKK1, DKK3, CER1, and NOTUM. It is possible that these secreted inhibitors protect the overlying epiblast from WNT signaling, thus preventing its differentiation. This is consistent with the role of these inhibitors *in vivo,* where Foxa2+ endoderm cells suppress Wnt signaling by secretion of inhibitors such as Cer1, Dkk1, and Sfrp1/5 (Scheibner et al., 2021).

Our study revealed that in pluripotent cells, WNT signaling induces posterior fates in a concentration-dependent manner only when NODAL signaling is inhibited, a previously unappreciated feature of the interaction between these pathways. Previous studies have uncovered complex molecular interactions in hPSCs. A direct interaction with SMAD2/3 can enhance ß-catenin’s transcriptional activity in the presence of both WNT and ACTIVIN, while during neural crest induction it is repressed by SMAD2/3 (Funa et al., 2015). When PI3K/ Akt signaling is active, Activin/Nodal signaling promotes self-renewal and pluripotency, while when PI3K is suppressed, Activin/Nodal synergizes with ß-catenin to induce mesendoderm differentiation (Singh et al., 2012). Moreover, it has also been shown that exposure of hPSCs to ACTIVIN activates SMAD2/3 but doesn’t induce differentiation. However, if hPSCs are exposed to WNT before ACTIVIN treatment, then ACTIVIN induces mesendodermal fates (Yoney et al., 2018; Yoney et al., 2022). Our previous work has shown that in pluripotent conditions, cells display an adaptive response to WNT signaling, in contrast to PS-like inducing conditions, where WNT signaling results in a sustained response, highlighting that WNT signaling dynamics are context specific (Camacho-Aguilar et al., 2023; Massey et al., 2019). Thus, complex molecular mechanisms regulate the interplay between WNT and NODAL signaling, and more work will be required to understand the specific mechanisms by which NODAL inhibition renders cells capable of posteriorizing in response to WNT, as found here.

In contrast, CHIR, the most commonly used Wnt activating reagent for inducing posterior cell fates *in vitro*, acts in a concentration-dependent manner even without Nodal inhibition. (Gouti et al., 2014; Lippman et al., 2015; Amin et al., 2016; Gouti et al., 2017; Metzis et al., 2018; Beccari et al., 2018; Matsuda et al., 2020; Veenvliet et al., 2020; Xu et al., 2021; Budjan et al., 2022; Miao et al., 2022; Sanaki-Matsumiya et al., 2022; Rito et al., 2023; García et al., 2023). Protocols that incorporate a pulse of CHIR with or without TGF-ß superfamily inhibition in three dimensions generate elongating organoids that model the posterior of the embryo including tailbud progenitors and their progeny (Beccari et al., 2018; van den Brink and van Oudenaarden, 2021; Sanaki-Matsumiya et al., 2022; Miao et al., 2023; Matsuda et al., 2020; Budjan et al., 2022). In this tailbud-like region, Sox2 and Bra co-expression is observed, while cells outside this region go on to express Tbx6 and markers of somitic mesoderm. In two dimensions, when WNT activation is combined with NODAL inhibition, we find similar patterns with SOX2+ and BRA+ cells surrounded by TBX6+ cells (Figure 5D,E), it would be interesting to test if this condition would be able to substitute the Wnt activation by CHIR in these organoid protocols.

While this manuscript was in preparation, recent work from other labs also created micropatterned models for cell fate decisions that occur downstream of Wnt signaling, but used CHIR in place of WNT. Consistent with our work here, these models also show that inhibition of Nodal signaling can increase PSM differentiation, and additionally found that properly timed inhibition can result in the generation of notochord cells (Rito et al., 2023; García et al., 2023). Interestingly, these systems did not undergo the complex morphogenesis that we observed, while here we did not observe a substantial number of cells with a notochord expression profile (FOXA2+,BRA+), even when stimulating with similar dynamics. These differences likely result from using CHIR as opposed to WNT and it would be interesting to better understand the mechanism behind this in the future. Consistently, Martyn et al. also observed different patterns of BRA expression between chemical activation (BMP4 + CHIR) and ligand (BMP4 + WNT3A) in micropatterned ECAD-KO colonies (Martyn et al., 2019B). Interestingly, in our studies, we obtained similar patterning either with CHIR alone or by combining WNT ligands with NODAL inhibition at 24 hours (Figure 6A,7A). Dynamic measurements of signaling in these cases show that similar fate patterns can occur with widely diverging signaling dynamics. Finally, we note that our findings do not challenge the validity of using CHIR for inducing cell fates. Instead, they complement existing systems and offer insights to consider when modeling development.

Our data, consistent with previous work, collectively supports that WNT and NODAL signaling are crucial for PS formation, and early disruption to either of these signals results in a general reduction in the expression of mesoderm and endoderm markers (Figure 6C). Our results suggest a model where, *in vivo,* initially, WNT induces NODAL, and these signals together induce primitive streak and subsequently combine to drive DE differentiation. As the embryo grows, the tailed region is reduced in NODAL signaling, allowing PSM differentiation. Future *in vivo* studies will be required to confirm and further elucidate these dynamics.

## Acknowledgments

We would like to thank Idse Heemskerk, Rosa Uribe, Hervé Turlier, and James Briscoe for helpful discussions on the project. All members of the Warmflash lab, past and present, for their helpful feedback and technical assistance across the development of this project. This work was supported by the National Science Foundation (MCB-2135296), the National Institute of General Medical Sciences (R01GM126122 and R35GM149328), and CONACYT (Consejo Nacional de Ciencia y Tecnología. México).

## Material and Methods

### Cell lines

Human embryonic stem cell line ESI-017 was used in this study and purchased from ESIBIO. Genomic and Karyotype analysis was performed by the manufacturer. GFP::ßcatenin, SOX2::mCitrine, ECAD::mCitrine, NODAL::mCitrine and NODAL Knockout cells were previously generated and characterized in this lab (Liu et al. 2022; Camacho-Aguilar et al., 2022 preprint). RUES2 cell line was a gift from Dr. Ali Brivanlou (Rockefeller University) and H9 cell line was a gift from Dr. Jianping Fu (University of Michigan). Pluripotency markers were regularly checked during and prior this study. All cell lines were tested regularly for mycoplasma and found negative.

### Routine cell culture

hESCs were routinely cultured in mTeSR1 (STEMCELL Technologies) medium, in Matrigel (Corning, 1:200 with DMEMF12) coated culture dishes at 37°c, 5% CO2. SOX2-mCitrine, ECAD-mCitrine, Nodal-mCitrine and Nodal Knockout cell lines were cultured in mTeSR Plus (STEMCELL Technologies). All experiments were performed with cells of passage no later than 50. Cells were routinely passaged with dispase (Fisher Scientific). Accutase was used for single cell suspension and seeding. ROCK-inhibitor (Ri) Y27672 (10 µM; Stem Cell Technologies) was always employed in single cell suspension conditions to improve viability. Blasticidin (5 µg/ ml) was used to maintain the fluorescent H2B transgene in the GFP::ßcatenin and SOX2::mCitrine cell lines. Puromycin (5 µg /ml) was to maintain the CFP::H2B transgene in the Nodal::mCitrine cell line.

### Standard culture experiments

ESI017 hESCs were washed with PBS first, then single-cell suspended with accutase for 5 min at 37°c. Collected cells were then centrifuged at 1000 rpm for 4 min and seeded on laminin-521 (Biolamina) coated 8-well ibidi dishes at a density of 40,000 cells/cm^2^ in mTeSR or mTeSR plus medium with Ri (10µM) for one hour. Medium removal was followed by two PBS washes. Induction medium was placed as indicated in Figure 1. N2B27 medium was prepared using a mixture of 48 ml of DMEM F12-Hamm’s 50:50 (Corning), 48 ml of Neurobasal (Life Technologies), 0.5 ml of N2 supplement (ThermoFisher), 1 ml of B27 supplement without vitamin A (ThermoFisher), 0.5 ml of Glutamax (Gibco) and 50 µl of ß-mercaptoethanol (Gibco). Induction medium (Wnt 1x) consisted of N2B27 with 200 ng/ml of FGF8 (Fisher Scientific) and 100 ng/ml WNT3A (Amsbio). Wnt 10x medium consisted of 200 ng/ml of FGF8 and 1000 ng/ml of WNT3A (Amsbio). All experiments were replenished with fresh medium daily.

### Micropatterned experiments

For all micropatterned experiments, glass CYTOO coverslips or 96 well CYTOO plates were coated with 5µg/ml Laminin-521 (Biolamina) in PBS +/+ (calcium and magnesium) for 3 hours at 37°c. Next, 1/3 of the total volume (for a well or a dish with a coverslip) was added with PBS +/+ gently. Then, 1/3 of the total volume is removed (the same volume that was added with PBS +/+ in the previous step); this is considered a serial wash. To remove laminin completely, 8 serial washes were performed, never allowing the wells or coverslips to dry. Plates were coated and used on the same day. Coverslips were either used on the same day or stored for one day at 4°c in PBS+/+.

For seeding micropatterns, hESCs were washed with PBS −/− once and suspended as single cells with accutase for 5 min at 37°C. Cells were collected, centrifuged at 1000 rpm for 4 minutes and resuspended in mTeSR or mTeSR Plus with Ri (10 µM). For CYTOO coverslips in 35mm dishes, 1.3 million cells were seeded and incubated for 1.5 hours at 37 °C. For CYTOO 96 well plates, 100,000 cells/well were seeded in 200 µl of medium and incubated for 1 hour at 37°C. For coverslips, two PBS +/+ serial washes and two full PBS +/+ washes were employed to remove unspecific binding. A full wash consists of full removal of volume and full addition of volume. For plates, one serial PBS +/+ wash and a full PBS +/+ wash were employed. Cells were then immediately treated with the induction medium indicated in the text and medium was refreshed daily. TGF-ß inhibitors consisted of LDN193189 (100 nM; Fisher Scientific) and SB431542 (10µM; Fisher Scientific). Chemical WNT activator employed was CHIR99021(10µM; Medchem Express).

### Immunostaining

All samples were fixed with 4%PFA for 20 min at room temperature (RT), followed by two PBS −/− washes. For standard culture immunostaining was performed as previously described (Nemashkalo et al., 2017). For micropatterns, blocking buffer was applied overnight (ON) at 4°C. Blocking buffer consists of 1% Triton-X (SIGMA), 10% DMSO (Life Technologies), and 5% Donkey serum in PBS−/−. Primary antibodies were incubated ON at 4°C in blocking solution, followed by two washes with washing solution of 30 min each. Next, secondary antibodies were incubated ON followed by six washes with washing solution of 30 min each. Washing solution is 0.1% Tween-20 (Sigma) in PBS. DAPI was added to the secondary antibody solution. Donkey Alexa Fluor (Life Technologies) antibodies were used for secondary staining in a 1:500 dilution. Coverslips were mounted in Fluoromount-G (Southern Biotech) and dried ON.

### Live cell imaging

GFP::ßcatenin, SOX2::mCitrine, ECAD::mCitrine, and NODAL::mCitrine were seeded as previously described, either on Cytoo plates or on Cytoo coverslips inside a CYTOO chip holder, treated as indicated, and then placed on the microscope for imaging. Three to five colonies of 500µm or 700µm were imaged per condition with a 20x NA 0.75 objective on an Olympus/Andor spinning disk confocal microscope. All acquisition were done at 37°C and 5% CO2 to maintain consistency. Medium replenishment was done each day in a laminar flow hood and samples were then returned to the microscope.

### Fixed cell imaging

Standard culture images were acquired with a 10x NA 0.4 objective on an Olympus IX83 inverted epifluorescent microscope. Micropatterns were imaged with a 20x NA 0.75 objective on an Olympus/Andor spinning disk confocal microscope. Micropatterns from supplemental figure S4 were acquired with an Olympus FV3000 confocal microscope with a 20x NA 0.75 objective.

### Quantification and analyses

All experiments were performed at least twice with consistent results. At least 5 colonies were analyzed per condition per experiment. Results in figures were calculated with all experiments. Micropattern’s images taken with the 20x objective were assembled using FIJI-pairwise stitching function. Ilastik was used for creating masks and 3D segmentation/classification. Mean Intensity was calculated as the average immunofluorescence intensity of the indicated marker in all quantified cells. Analysis and graphs were done through custom Matlab written code https://github.com/warmflashlab/MAOS-WNTNODAL.

### Cell sorting

The SOX2::mCitrine cell line was seeded onto three CYTOO coverslips and treated as indicated. Three coverslips per condition were utilized in order to generate enough cells for sorting. Cells were single-cell suspended with Accutase at 37°C for 5 minutes. Cells were collected, centrifuged at 1000 rpm for 4 minutes and re-suspended in PBS-EDTA (10 mM). Cells were then filtered through a 5ml Polystyrene Round-Bottom tube with cell-strainer mesh 40um cap (FALCON). A Sony SH800S Cell Sorter with a 488 nm laser was used for separating the SOX2 positive cells from the negative ones.

### RNA extraction

RNA extraction was performed with Dneasy Blood and Tissue Kit (Qiagen).

### Bulk RNA sequencing data pre-processing

To quantify the abundance of transcripts for each sample, Salmon v1.9.01 was used. Reads were aligned to the human transcriptome (GRCh38) index for salmon, with salmon index using the selective alignment method (salmon_sa_index:default in http://refgenomes.databio.org/v3/ genomes/splash/2230c535660fb4774114bfa966a62f823fdb6d21acf138d4). We quantified transcripts with Salmon using the -l A flag to infer the library type (paired-end) automatically and the –validateMappings flag to use the selective alignment method. The resulting datasets were then processed using the tximeta2 package in R, and the counts matrix was exported for downstream analyses.

### Bulk RNA sequencing data integration with human and monkey single-cell RNA sequencing databases

To analyze the identity of the in vitro cultured cells, we integrated our bulk RNA sequencing data with reference in vivo human single-cell transcriptomics dataset (Tyser et al., 2021). The reference dataset was first pre-processed as described in Tyser et al. using scanpy v1.9.15 (Wolf, Angerer and Theis, 2018). We kept cells with more than 2,000 genes detected, and genes that have at least 1,000 total counts using the filter_cells and filter_genes functions. We then kept the top 4,000 highly variable genes (HVGs) using the high_variable_genes function, resulting in a total of 1,195 cells. We then processed our bulk RNA sequencing dataset for integration. We combined the transcript-level counts to obtain the corresponding gene-level counts. Then, we merged the Annotated Data objects of our bulk RNA seq dataset and the processed human Annotated Data objects using the concatenate function using the option “inner” for the join parameter to intersect the genes present in each object. This resulted in one merged Annotated Data object, with the human reference data, with a total of 3,649 genes. To account for batch effects in the downstream analysis of the merged datasets, we added a batch label, where cells coming from the reference Annotated Data object were labeled as batch 0, while the two repeats of the bulk dataset were labeled as batches 1 and 2, respectively. We then applied the scvi.model.SCVI6 function in each merged Annotated Data object separately to obtain a representation that integrated our bulk RNA sequencing data and the reference human dataset. We chose “counts” as the layer option, we took into account the batch labels in the batch_key option for the integration, and used two hidden layers (n_layers=2) with 30 as the dimension of the latent space (n_latent=30). Using the resulting representation of the data, we then computed a neighborhood graph of the data using the neighbors function from scanpy with default settings, the result of which was used to obtain a UMAP plot of the data (using the tl.umap function with default parameters). To identify clusters, we applied the Leiden algorithm with a resolution of 0.7.

## Notes

### Competing Interest Statement

AW is a cofounder and holds equity in Simbryo Technologies. The work presented here is not related to the interests of this commercial entity. All other authors declare no competing interests.

