## Supplementary figures for "Endogenous Nodal switches Wnt interpretation from posteriorization to germ layer differentiation in geometrically constrained human pluripotent cells"

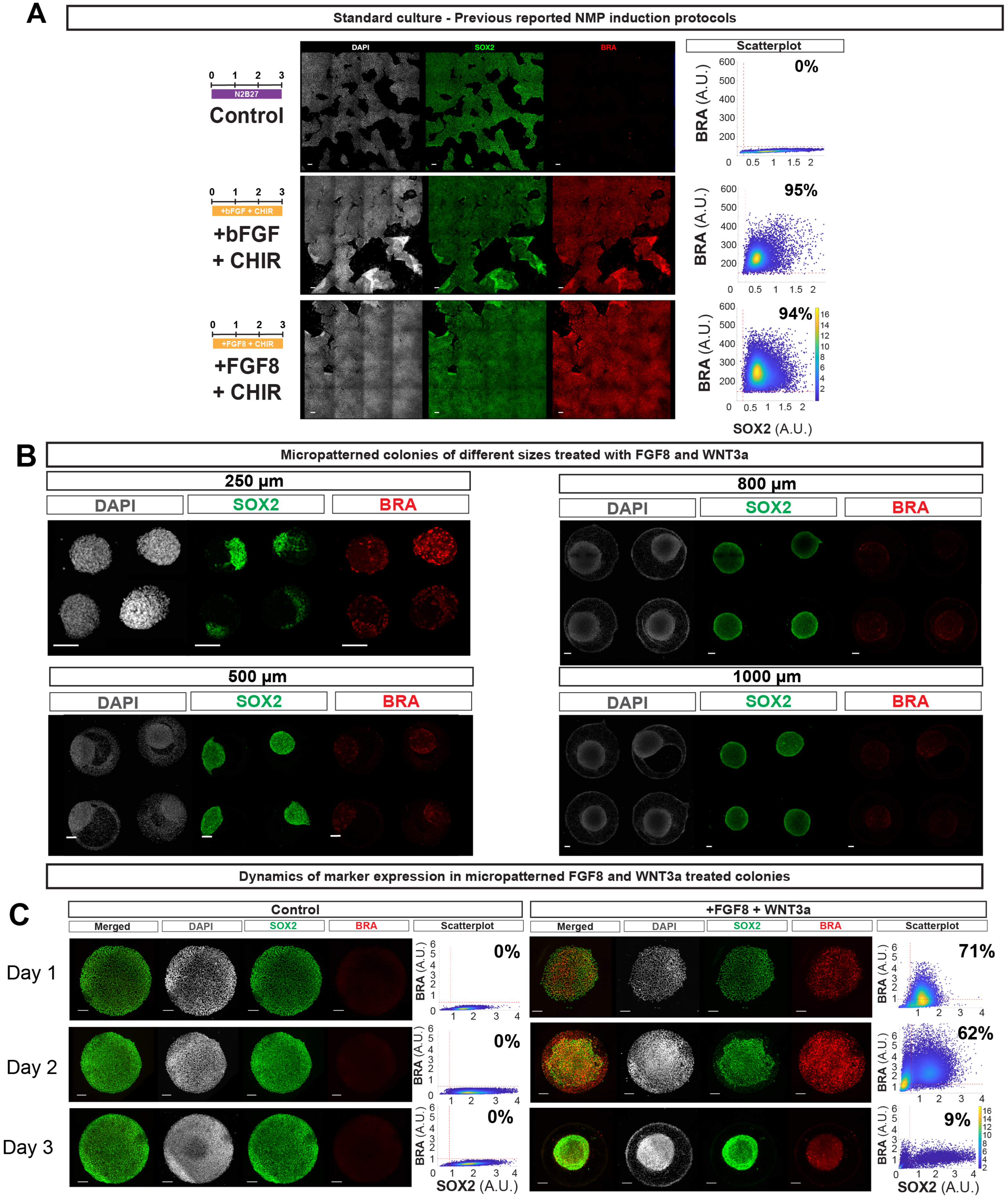


**Figure S1. NMP induction in standard culture.**

(A) Schematics (left), representative images (center) and quantifications (right) of cells subjected to NMP protocols in standard culture. Protocols are as in Gouti et al and Lippmann et al. (B) Representative immunofluorescent images of micropatterned colonies of the indicated sizes (C) Representative immunofluorescent images and quantifications of micropatterned colonies from day 1 to end of treatment. In scatterplots, each dot represents a cell, red lines represent the 99th-percent quantiles of the distributions obtained from the negative control for SOX2 and BRA, and the color bar represents the probability density $1x10^{-7}$. Number in the top right indicates the percentage of cells expressing both SOX2 and BRA markers. All images are max projections. All scale bars are 100µm.

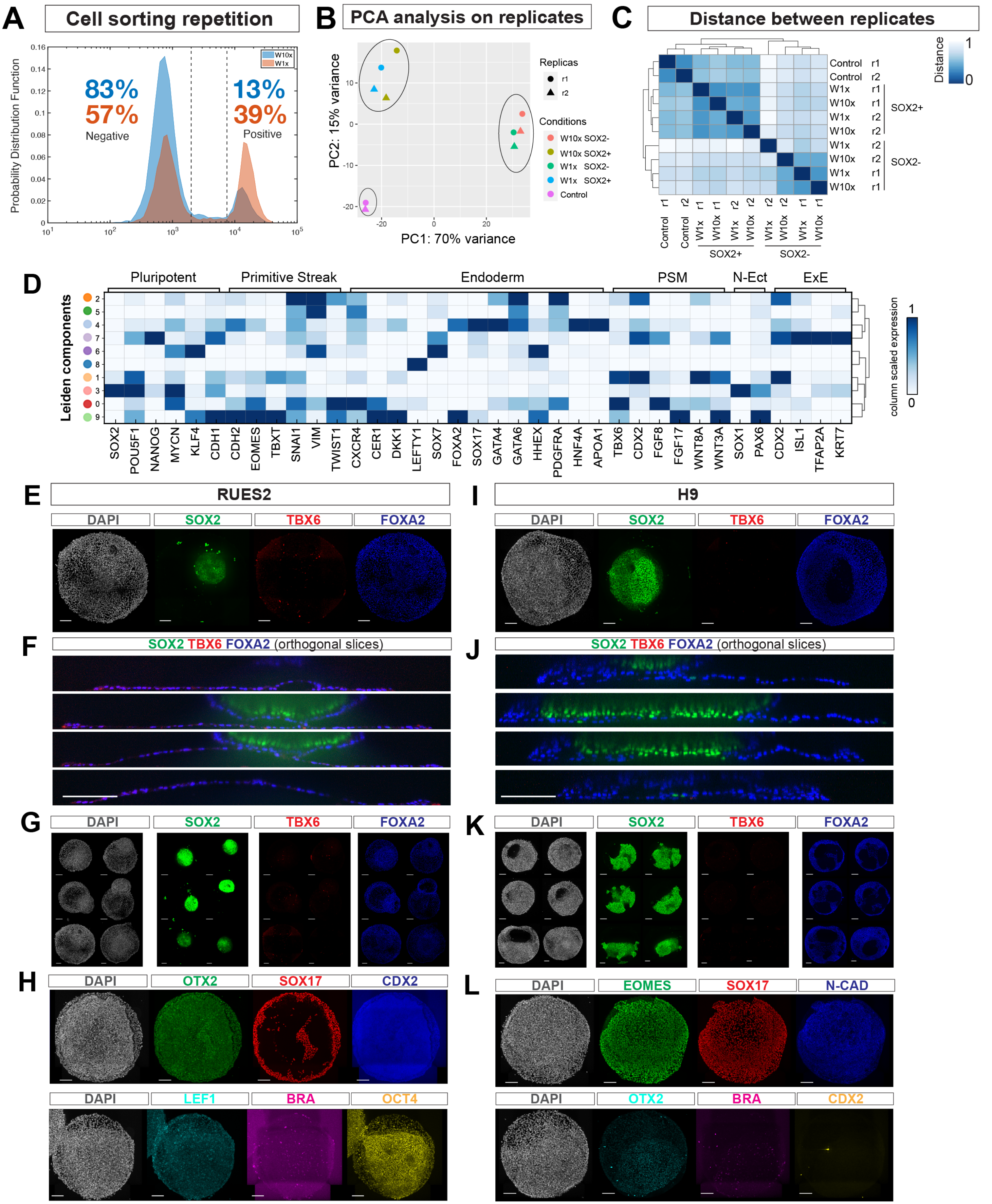


**Figure S2. Reproducibility of WNT3A and FGF8 treatment on micropatterns in replicates of RNA-sequences and for different cell lines.**
(A) Probability distribution function histogram of the repeated cell sorting experiment. (B) Principal component analysis of gene expression data from RNA sequencing. (C) Correlation heatmap between experimental replicates. (D) Heatmap showing mean RNA levels for the indicated representative genes in each Leiden component. PSM - Presomitic mesoderm, N-Ect - Neural ectoderm, ExE - Extraembryonic ectoderm. (E-H) Representative images from the RUES2 cell line treated with WNT3A and FGF8. (E) 800 µm colony (F) Orthogonal slices of colony (E). (G) 500 µm colonies (H) 700 µm colonies. (I-L) Representative images of H9 cells treated with WNT3A and FGF8. (I) 800 µm colony chip. (J) Orthogonal slices of colony (K). (J) 500 µm colonies (L) 700 µm colonies. Images in (E,I,G,K, H and L) are max projections. All scale bars are 100µm. (E-G, I-K) are performed on Cytoo coverslips, (H,L) are performed on Cytoo 96 well plates.

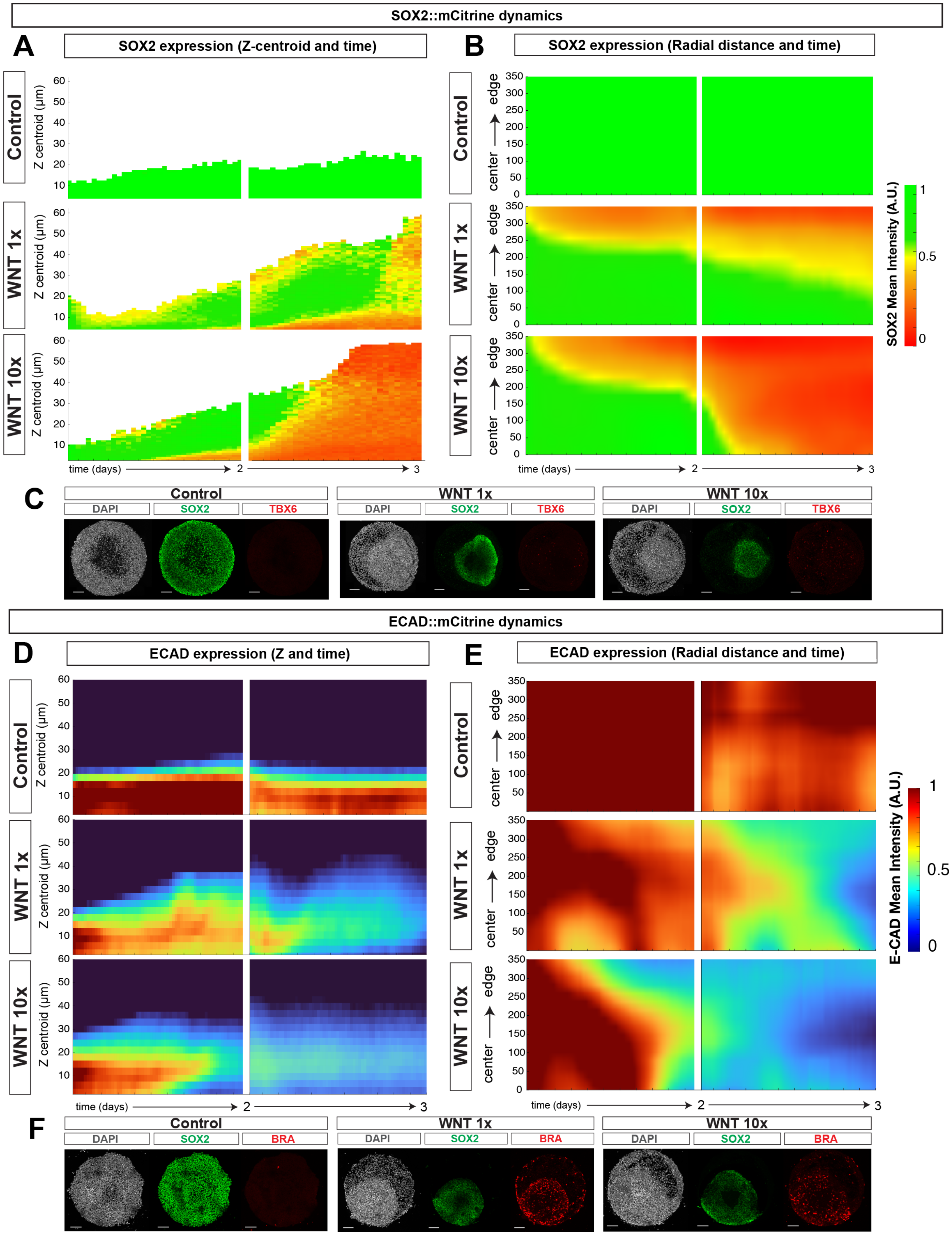


**Figure S3. SOX2::mCitrine and ECAD::mCitrine dynamics for day 2 and 3 of treatment.**
(A,B) Kymographs quantifying nuclear SOX2 mean intensity levels as a function of Z-centroid and time (A) or as a function of distance from the center of the colony and. time (B). (C) Representative immunofluorescent images of the SOX2::mCitrine cell line immunostained at end of treatment. (D, E) Kymograph quantifying E-CAD mean intensity levels as a function of Z and time (D) or as a function of distance from the center of the colony and time.(F) Representative immunofluorescent images of the ECAD::mCitrine cell line immunostained at the end of treatment. All images are max projections. Scale bars are 100μm. In (A,B,D,E), each colored square represents the mean intensity of the reporter for all cells quantified at that position during that time.

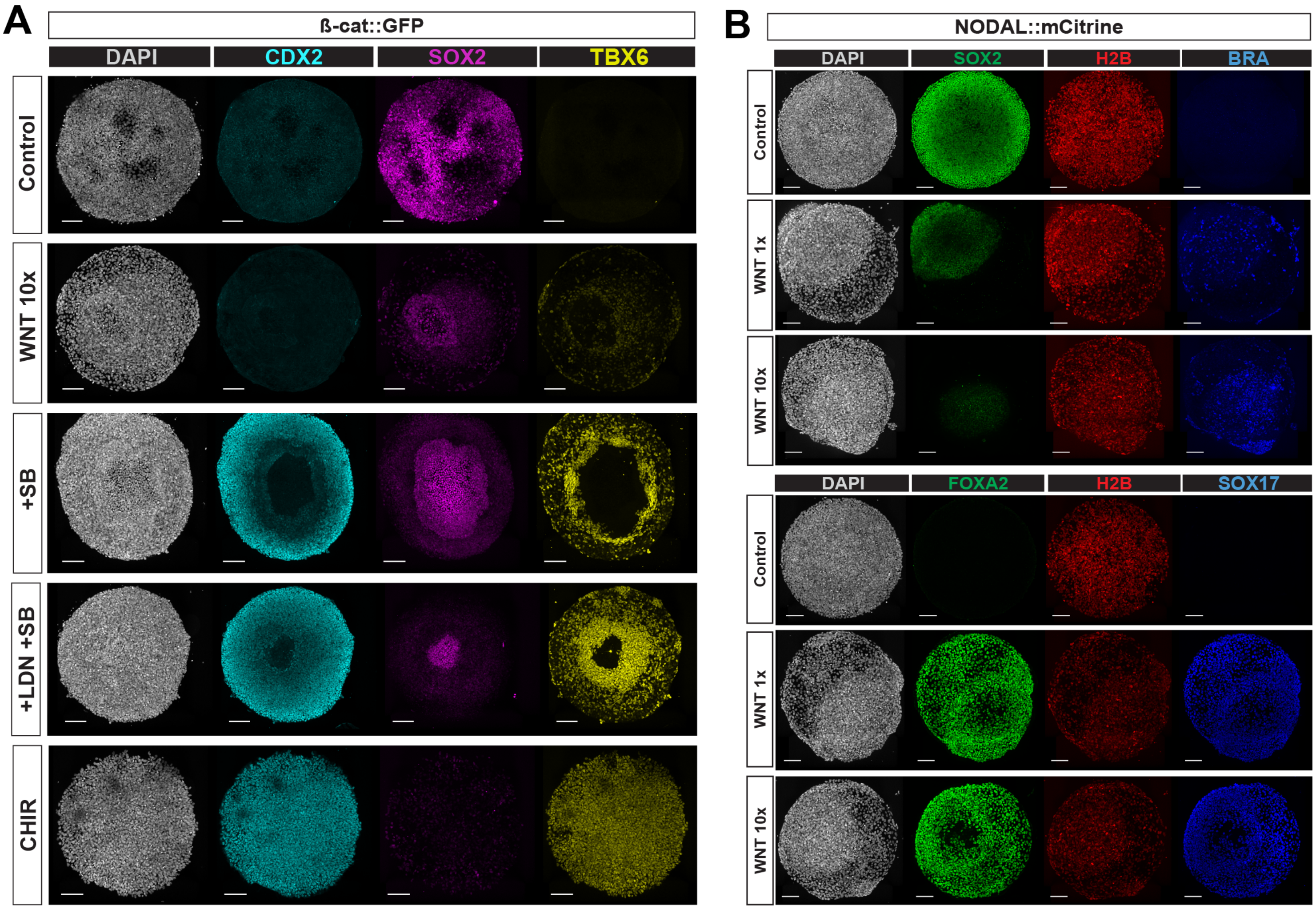


**Figure S4. Signaling reporter cells that have undergone live-cell imaging display the same patterning as WT hESCs.**
(A) Representative immunofluorescent images of ßcat::GFP cell line under different conditions after live imaging.
(C) Representative immunofluorescent images of NODAL::mCitrine cell line of 700μm under different conditions after live imaging.
All images are max projections. Scale bars are 100 μm.

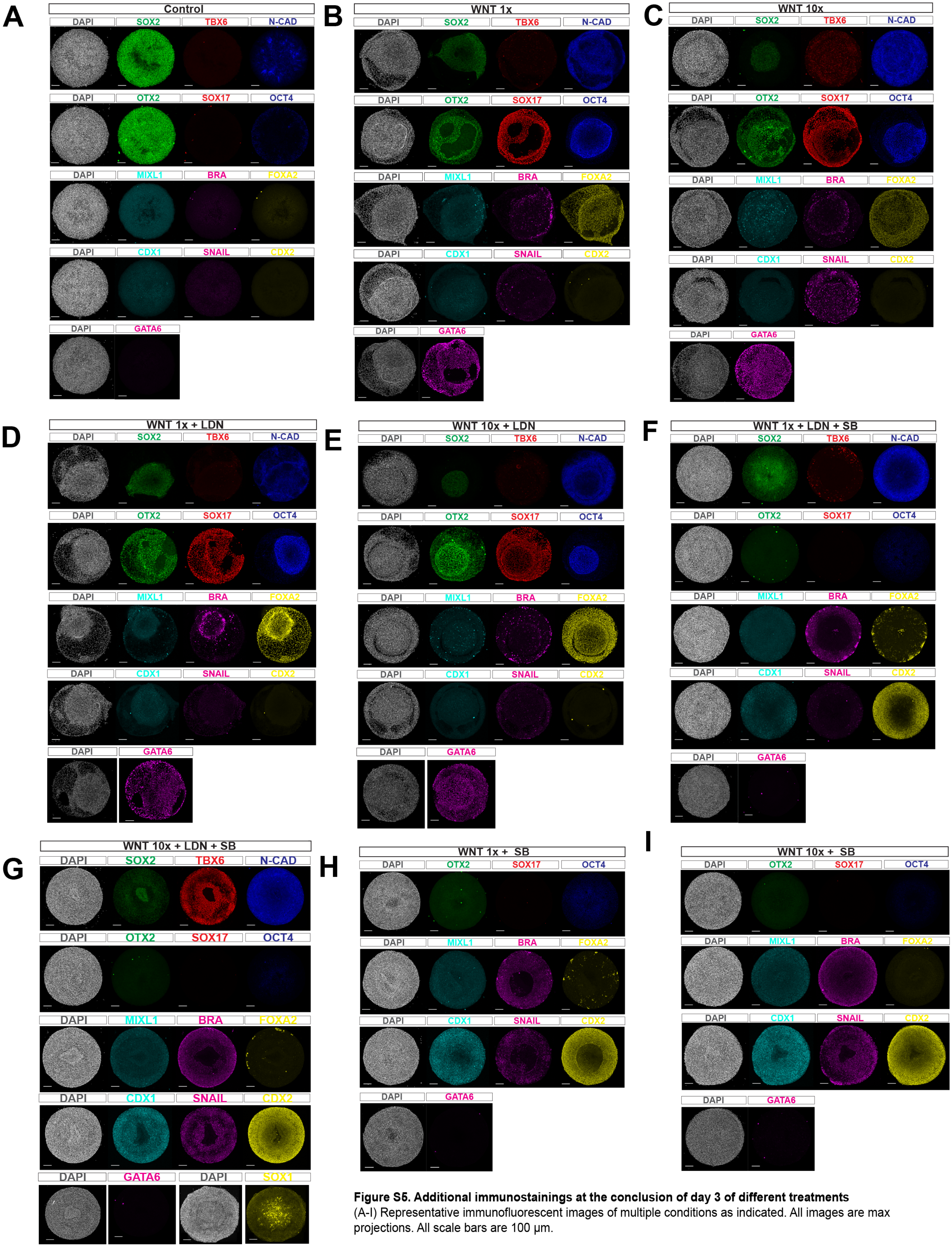


**Figure S5. Additional immunostainings at the conclusion of day 3 of different treatments**

(A-I) Representative immunofluorescent images of multiple conditions as indicated. All images are max projections. All scale bars are 100 μm.

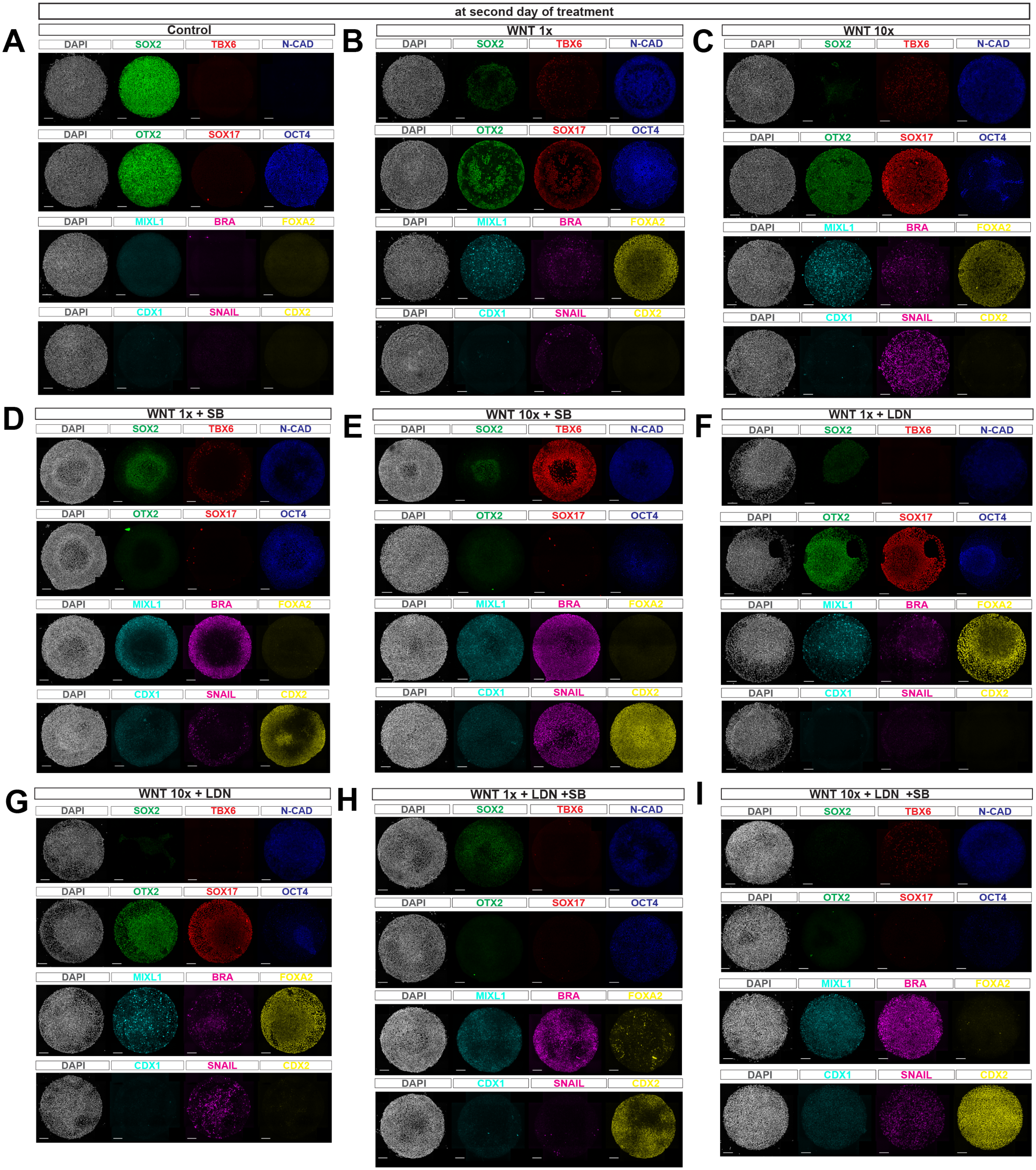


**Figure S6. Multiple immunostainings after day 2 of different treatments.**
(A-I) Representative immunofluorescent images of multiple conditions as indicated. All conditions were fixed at the end of day 2 of treatment. All images are max projections. All scale bars are 100 μm.

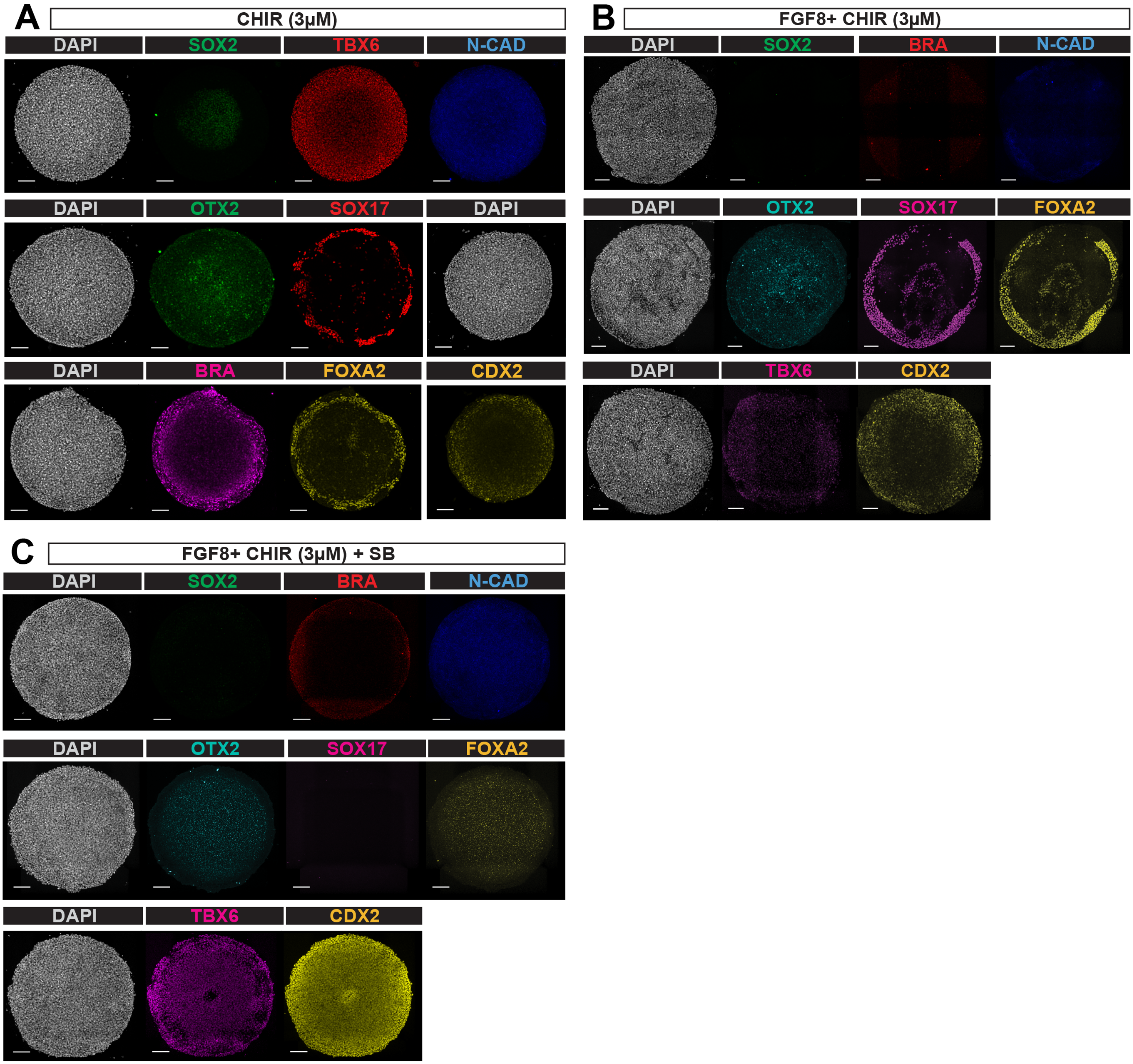


**Figure S7. Low CHIR elicits DE cell fates, with or without FGF8, unless TGF-ß is inhibited**
(A) Representative immunofluorescent images of colonies treated with 3μM of CHIR.
(B) Representative immunofluorescent images of colonies treated with 3μM of CHIR and FGF8 (200 ng/ml).
(C) Representative immunofluorescent images of colonies treated with 3μM of CHIR, FGF8 (200 ng/ml) and SB (10 μM). All images are max proejctions. All scale bars are 100 μm.

### Supplemental figures

**Figure S1. NMP induction in standard culture.**
(A) Schematics (left), representative images (center) and quantifications (right) of cells subjected to NMP protocols in standard culture. Protocols are as in Gouti et al and Lippmann et al. (B) Representative immunofluorescent images of micropatterned colonies of the indicated sizes (C) Representative immunofluorescent images and quantifications of micropatterned colonies from day 1 to end of treatment. In scatterplots, each dot represents a cell, red lines represent the 99th-percent quantiles of the distributions obtained from the negative control for SOX2 and BRA, and the color bar represents the probability density $1x10^{-7}$. Number in the top right indicates the percentage of cells expressing both SOX2 and BRA markers. All images are max projections. All images are max projections. All scale bars are 100μm.

**Figure S2. Reproducibility of WNT3A and FGF8 treatment on micropatterns in replicates of RNA-sequences and for different cell lines.**
(A) Probability distribution function histogram of the repeated cell sorting experiment. (B) Principal component analysis of gene expression data from RNA sequencing. (C) Correlation heatmap between experimental replicates. (D) Heatmap showing mean RNA levels for the indicated representative genes in each Leiden component. PSM - Presomitic mesoderm, N-Ect - Neural ectoderm, ExE - Extraembryonic ectoderm. (E-H) Representative images from the RUES2 cell line treated with WNT3A and FGF8. (E) 800 μm colony (F) Orthogonal slices of colony (E). (G) 500 μm colonies (H) 700 μm colonies. (I-L) Representative images of H9 cells treated with WNT3A and FGF8. (I) 800 μm colony chip. (J) Orthogonal slices of colony (K). (J) 500 μm colonies (L) 700 μm colonies. Images in (E,I,G,K, H and L) are max projections. All scale bars are 100μm. (E-G, I-K) are performed on Cytoo coverslips, (H,L) are performed on Cytoo 96 well plates.

**Figure S3. SOX2::mCitrine and ECAD::mCitrine dynamics for day 2 and 3 of treatment.**
(A,B) Kymographs quantifying nuclear SOX2 mean intensity levels as a function of Z-centroid and time (A) or as a function of distance from the center of the colony and. time (B). (C) Representative immunofluorescent images of the SOX2::mCitrine cell line immunostained at end of treatment. (D, E) Kymograph quantifying E-CAD mean intensity levels as a function of Z-centroid and time (D) or as a function of distance from the center of the colony and time.(F) Representative immunofluorescent images of the ECAD::mCitrine cell line immunostained at the end of treatment. All images are max projections. Scale bars are 100μm. In (A,B,D,E), each colored square represents the mean intensity of the reporter for all cells quantified at that position during that time.

**Figure S4. Signaling reporter cells that have undergone live-cell imaging display the same patterning as WT hESCs.**
(A) Representative immunofluorescent images of ßcat::GFP cell line under different conditions after live imaging. (C) Representative immunofluorescent images of NODAL::mCitrine cell line of 700μm under different conditions after live imaging. All images are max projections. Scale bars are 100 μm.

**Figure S5. Additional immunostainings at the conclusion of day 3 of different treatments**
(A-I) Representative immunofluorescent images of multiple conditions as indicated. All images are max projections. All scale bars are 100 μm.

**Figure S6. Multiple immunostainings after day 2 of different treatments.**
(A-I) Representative immunofluorescent images of multiple conditions as indicated. All conditions were fixed at the end of day 2 of treatment. All images are max projections. All scale bars are 100 μm.

**Figure S7. Low CHIR elicits DE cell fates, with or without FGF8, unless TGF-ß is inhibited**
(A) Representative immunofluorescent images of colonies treated with 3μM of CHIR.
(B) Representative immunofluorescent images of colonies treated with 3μM of CHIR and FGF8 (200 ng/ml). (C) Representative immunofluorescent images of colonies treated with 3μM of CHIR, FGF8 (200 ng/ml) and SB (10 μM). All images are max projections. All scale bars are 100 μm.

**Movie S1. 3D rendering of 500um diameter colony treated with WNT3A and FGF8 for three days.** DAPI (blue), SOX2 (green), SOX17(red).

**Movie S2. Live imaging showing maximum intensity projections of a colony of the SOX2::mCitrine reporter.** z-stacks were taken every 40 minutes from 24 to 72 hours after treatment with WNT3A and FGF8. Colony is 700 μm diameter. H2B (red), SOX2 (green). Time stamp is in hours:minutes.

**Movie S3. Live imaging showing maximum intensity projections of a colony of the ECAD::mCitrine reporter.** z-stacks were taken every 40 minutes from 24 to 72 hours after treatment with WNT3A and FGF8. Colony is 700 μm diameter. H2B (red), SOX2 (green). Time stamp is in hours:minutes.
